# Deep5hmC: Predicting genome-wide 5-Hydroxymethylcytosine landscape via a multimodal deep learning model

**DOI:** 10.1101/2024.03.04.583444

**Authors:** Xin Ma, Sai Ritesh Thela, Fengdi Zhao, Bing Yao, Zhexing Wen, Peng Jin, Jinying Zhao, Li Chen

## Abstract

5-hydroxymethylcytosine (5hmC), a critical epigenetic mark with a significant role in regulating tissue-specific gene expression, is essential for understanding the dynamic functions of the human genome. Using tissue-specific 5hmC sequencing data, we introduce Deep5hmC, a multimodal deep learning framework that integrates both the DNA sequence and the histone modification information to predict genome-wide 5hmC modification. The multimodal design of Deep5hmC demonstrates remarkable improvement in predicting both qualitative and quantitative 5hmC modification compared to unimodal versions of Deep5hmC and state-of-the-art machine learning methods. This improvement is demonstrated through benchmarking on a comprehensive set of 5hmC sequencing data collected at four time points during forebrain organoid development and across 17 human tissues. Notably, Deep5hmC showcases its practical utility by accurately predicting gene expression and identifying differentially hydroxymethylated regions in a case-control study of Alzheimer’s disease.

## Introduction

5-hydroxymethylcytosine (5hmC) modification is one important intermediate state among a successive of states in active demethylation, which includes 5-Methylcytosine (5mC), 5hmC, 5-formylcytosine (5fC), and 5-carboxylcytosine (5caC). The generation of 5hmC occurs through the oxidation of 5mC by the ten-eleven translocation (TET) protein family, and it is specifically recognized by 5hmC-binding proteins (Spruijt et al., 2013; Tahiliani et al., 2009). In the nervous system, 5hmC plays a critical role in neurodevelopment and neurological function. It has been found to be enriched in embryonic stem cells and neuronal cells, regulating neuronal-specific gene expression during neural progenitor cell differentiation (Kriaucionis & Heintz, 2009; X. Li et al., 2017). Abnormalities in 5hmC distribution and enrichment can be critical factors contributing to neurodegenerative diseases such as Huntington’s disease, Autism spectrum disorder and Alzheimer’s disease (AD) (Bernstein et al., 2016; Cheng et al., 2018; Coppieters et al., 2014; Kuehner et al., 2021; Qin et al., 2020; Wang et al., 2013). Beyond neurodegenerative diseases, 5hmC also plays a significant role in cancer development and treatment. Genome-wide mapping of 5hmC reveals that loss of 5hmC is an epigenetic hallmark of melanoma and medulloblastoma (Lian et al., 2012; Stahl et al., 2021; Zhao et al., 2021). Additionally, 5hmC in circulating cell-free DNA serves as diagnostic biomarkers for colorectal, gastric, pancreatic cancer, acute myeloid leukemia and other common human cancer types (Gao et al., 2019; W. Li et al., 2017; Shao et al., 2022). Importantly, 5hmC-based biomarkers of circulating cfDNA have demonstrated high predictiveness of cancer stage and superior to conventional biomarkers (Guler et al., 2020; W. Li et al., 2017; Song et al., 2017).

The emergence of next generation sequencing has facilitated the genome-wide profiling of 5hmc modification. Among 5hmC sequencing technologies, antibody-based immunoprecipitation and sequencing of hydroxymethylated DNA (hMeDIP-seq) (Weber et al., 2005) as well as 5hmC-selective chemical labeling method (e.g. 5hmC-Seal) (Han et al., 2016; Song et al., 2011), have become cost-effective methods to map genome-wide 5hmC signals. These methods aim to capture and enrich 5hmC methylated DNA fragments, followed by next-generation sequencing. Similar to ChIP-seq analysis, the enriched region of 5hmC, deemed as “peak”, can be identified by peak-calling algorithm such as MACS (Zhang et al., 2008). Due to their popularity, hMeDIP-seq/5hmC-Seal and other similar protocols have been widely adopted to explore the distribution and patterns of genome-wide 5hmC in various tissues, cell types (Cui et al., 2020; W. Li et al., 2017) and diseases (Bernstein et al., 2016; Cheng et al., 2018; Kuehner et al., 2021; Qin et al., 2020). In addition, 5hmC sequencing aids in investigating the association between 5hmC and other genomic elements. For instance, 5hmC has been found to co-localize with gene bodies and enhancers known to activate gene expression, and it is positively correlated with gene expression (He et al., 2021). Moreover, 5hmC is significantly enriched in histone modifications associated with active enhancers, such as H3K4me1 and H3K27ac (Cui et al., 2020; Stroud et al., 2011). Furthermore, the distribution and enrichment of 5hmC exhibit tissue-specificity, evident in its preferential enrichment on tissue-specific gene bodies and enhancers (Li & Liu, 2011).

Nevertheless, it is still costly to conduct a deep sequencing for an accurate identification of 5hmC modification. In addition, 5hmC experiments may suffer from underpowering due to insufficient sequencing depth or various artifacts, leading to a limited detection of 5hmC modification sites. Moreover, 5hmC profiles exhibit dynamic changes across tissues and cell types. To overcome these challenges, computational models have been developed to enable an *in silico* genome-wide prediction of 5hmC profiles. The general principle of these methods is to treat DNA sequence within a genomic region as the model input and predict the probability of the region being a 5hmC peak. The key distinction between these approaches lies in the feature engineering applied to DNA sequences and successive use of machine learning algorithms. For example, iRNA5hmC-PS adopts k-mer (k=2,3) frequency and uses logistic regression for predicting the presence of 5hmC peaks (Ahmed et al., 2020). iRhm5CNN utilizes a simple convolution neural network, which employs one-hot encoding DNA sequence as the model input to predict the occurrence of 5hmC peaks (Ali et al., 2021). Given that the resolution of 5hmC peaks closely aligned with that of ChIP-seq peaks, deep learning methods designed for predicting the binding sites of transcriptional factor, histone modification sites and open chromatin regions such as DeepSEA (Zhou & Troyanskaya, 2015), which aims to predict epigenetic signals across hundreds of tissues and cell types in a multi-task framework, can be readily adapted for the task for predicting 5hmC peaks.

Despite the success of existing computational methods for predicting 5hmC modification, there are still challenges to be addressed. First, there is a lack of computational methods specifically designed for predicting tissue/cell type-specific 5hmC modification on DNA is lacking, and alternative methods substituted for this purpose such as DeepDEA may be suboptimal. Second, the relationship among 5hmC, histone modification and gene expression is rarely explored in the predictive modeling of 5hmC modification. Lastly, current methods primarily concentrate on classifying binary 5hmC peaks while overlooking the quantitative variation of 5hmC modification. To address these challenges, we introduce a novel multi-modal deep learning framework named Deep5hmC, which aims to enhance the prediction of tissue/cell type-specific genome-wide 5hmC modification by incorporating information from both DNA sequence and histone modification. The contribution of our work lies on following aspects: (i) Deep5hmC leverages both DNA sequence and histone modifications to improve the prediction of 5hmC modification in both qualitative (i.e., 5hmC peaks) and quantitative prediction (i.e., normalized 5hmC reads); (ii) Deep5hmC is developed and evaluated using a comprehensive set of 5hmC sequencing (5hmc-seq) data collected at four time points during forebrain organoid development and across 17 human tissues. The extensive dataset demonstrates the power of Deep5hmC in predicting tissue-specific 5hmC modification; (iii) Deep5hmC is further assessed using one 5hmC-seq data in one case-control study of Alzheimer’s disease to demonstrate its broad utility in predicting differentially hydroxymethylated regions (DhMR) within the context of the disease; (iv) an extension of Deep5hmC is to quantify gene expression by predicted quantitative 5hmC modification within gene bodies; (v) Deep5hmC is released as an open-source python toolkit, which can benefit the epigenetic research community. As a result, we demonstrate that inclusion of histone modification leads to improved prediction performance of Deep5hmC for both qualitative and quantitative 5hmC modification. Importantly, Deep5hmC outperforms competing machine learning approaches for the same purpose. In addition, Deep5hmC achieves an accurate prediction of gene expression and is also powerful for predicting DhMRs.

## Results

### Overview of Deep5hmC

The workflow of the deep learning framework, Deep5hmC is demonstrated in **Fig. 1**. The labelled training set can be derived from tissue/cell-type specific 5hmC-enriched region (i.e., peaks) in one condition (**Fig. 1A**) or DhMRs in a case-control study (**Fig. 1B**) (e.g., disease versus healthy control). Accordingly, one-hot encoding DNA sequence in peaks or DhMRs serves as the input for the sequence modality (**Fig 1C, D**). In addition, histone features are obtained from histone ChIP-seq from public consortiums such as ENCODE or Roadmap Epigenomics (**Fig. 1A**), with matched tissue/cell-type as the 5hmC-seq. Only the histone features in the neighborhoods of the 5hmC peaks/DhMRs are considered as the input for the histone modality. The sequence modality and histone modality each go through their own convolutional neural networks (CNN) to derive separate feature representations, which are later joined via the MFB fusion layer. The output of the MFB fusion layer further connects to fully connected layers and the output layer afterwards. Depending on the preparation of training set and prediction mission, the Deep5hmC consists of four modules, including Deep5hmC-binary, Deep5hmC-cont, Deep5hmC-gene and Deep5hmC-diff. Specifically, Deep5hmC-binary takes the labelled 5hmC peaks and non-peaks as the training set to identify the 5hmC-enriched regions (More details in section “Evaluating Deep5hmC for predicting binary 5hmC modification sites”). Deep5hmC-cont takes normalized read counts in 5hmC peaks and aim to predict the continuous 5hmC modification genome-wide (More details in section “Evaluating Deep5hmC for predicting continuous 5hmC modification”). By leveraging Deep5hmC-cont, Deep5hmC-gene aggregates the predictions of Deep5hmC-cont in the gene bodies as a surrogate for predicted gene expression (More details in section “Evaluating Deep5hmC in predicting gene expression”). Different from Deep5hmC-binary, Deep5hmC-diff takes the labelled DhMRs/non-DhMRs in a case-control design of 5hmC-seq as the training set, and derives histone features from a similar case-control design of histone ChIP-seq. Deep5hmC-diff aims to predict genome-wide DhMRs and discover *de novo* DhMRs that may be absent in the training set (More details in section “Evaluating Deep5hmC for predicting differential hydroxymethylated regions”). Overall, the four modules in the deep learning framework Deep5hmC will provide a comprehensive assessment of genome-wide tissue/cell type-specific 5hmC modification in either qualitative or quantitative manner as well as genome-wide DhMRs. In addition, it allows the prediction of gene expression using predicted 5hmC modification. The details and evaluation for each module of Deep5hmC will be elaborated in the subsequent sections.

**Figure 1.**
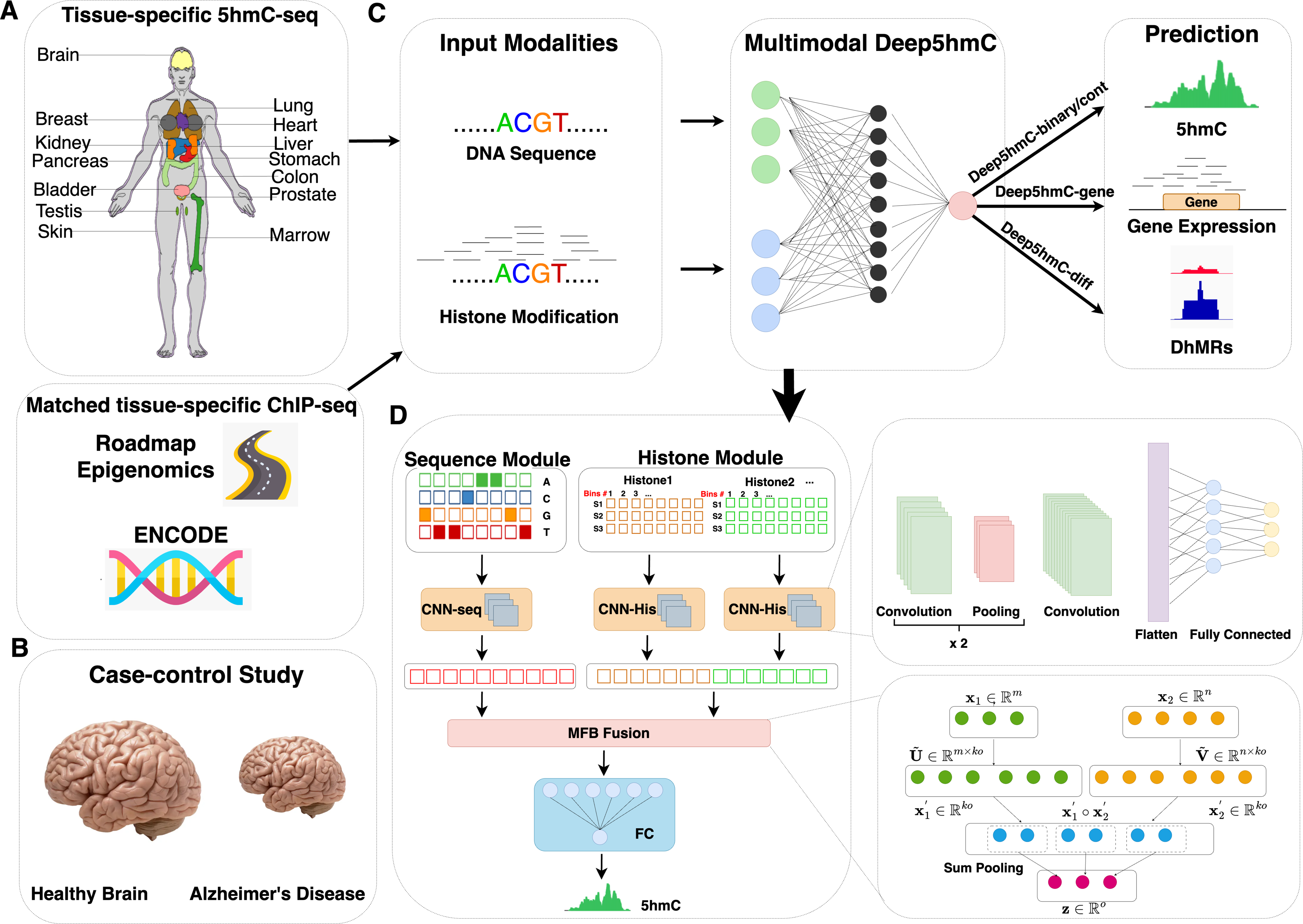
Overview of Deep5hmC. **A**. The training set of Deep5hmC can be derived from matched 5hmC-seq and histone ChIP-seq from one condition. Specifically, the 5hmC-seq data can be collected from tissue-specific human tissues, which include bladder, brain, breast, heart, kidney, liver, lung, marrow, ovary (female), pancreas, placenta (female), prostate (male), colon (sigmoid), colon (transverse), skin, stomach and testis (male). The matched tissue-specific histone ChIP-seq data are collected according from public consortiums such as Roadmap Epigenomics and ENCODE. In this context, Deep5hmC aims to predict genome-wide 5hmC modification in a single condition. **B**. The training set of Deep5hmC can also be derived from matched 5hmC-seq and histone ChIP-seq from a case-control study (e.g., Alzheimer’s Disease vs healthy control) for predicting differentially hydroxymethylated regions (DhMRs). **C.** Deep5hmC is a multimodal deep learning model to improve the prediction of tissue/cell type-specific genome-wide 5hmC modification by leveraging both DNA sequence and histone modification. Deep5hmC consists of four modules, including Deep5hmC-binary, Deep5hmC-cont, Deep5hmC-gene and Deep5hmC-diff. Specifically, Deep5hmC-binary takes the labelled 5hmC peaks and non-peaks as the training set to identify the 5hmC enriched regions. Deep5hmC-cont takes the normalized read counts in 5hmC peaks and aim to predict the continuous 5hmC modification genome-wide. By leveraging Deep5hmC-cont, Deep5hmC-gene aggregates the predictions of Deep5hmC-cont in the gene bodies as the surrogate for the predicted gene expression. Different from Deep5hmC-binary, Deep5hmC-diff takes the labelled DhMRs/non-DhMRs in a case-control design of 5hmC-seq as the training set to predict genome-wide DhMRs and may discover *de novo* DhMRs. **D.** Model architecture of Deep5hmC. Deep5hmC consists of both sequence modality and histone modality consisting of their own convolutional neural networks (CNN) to derive separate feature representations, which will be joined later via the Multi-modal Factorized Bilinear pooling (MFB) fusion layer. The output of the MFB fusion layer will further connect to fully connected layers and the output layer afterwards.

### Distribution pattern of histone modification around 5hmC peaks

We conduct a real data exploration by integrating tissue matched 5hmC-seq data and histone ChIP-seq data to evaluate the potential of histone modification as informative features for predicting 5hmC modification. Without loss of generality, we gather EB 5hmC peaks from “Forebrain Organoid” and ChIP-seq data in “Brain Angular Gyrus” involving seven histone marks from Roadmap Epigenomics (**Supplementary Table S1**). The seven histone marks consists of H3K4me1, H3K27ac and H3K9ac associated with active enhancers; H3K4me3 associated with active promoters; H3K36me3 associated with active expressed gene bodies; and repressive marks such as H3K9me3 and H3K27me3. To characterize the histone modification patterns around 5hmC modification sites, we acquire and average the histone features with dimensions of 1×41 in the neighborhood of each 5hmC peak for the positive and negative sets. The histone features represent essentially normalized 5hmC read counts, which are created by segmenting an extended genomic region of 10kb both upstream and downstream of each 5hmC peak into 41 1kb windows, with a sliding size 500bp and counting reads for each 1kb windows (More details in “Multimodal features”). Performing the Wilcoxon rank-sum test on the two sets of histone features for each histone mark, we find that the distribution of histone features is significantly different between positive and negative 5hmC peaks (pvalue<0.05) (**Fig. 2**). This observation suggests that histone marks are informative for predicting 5hmC modification.

**Figure 2.**
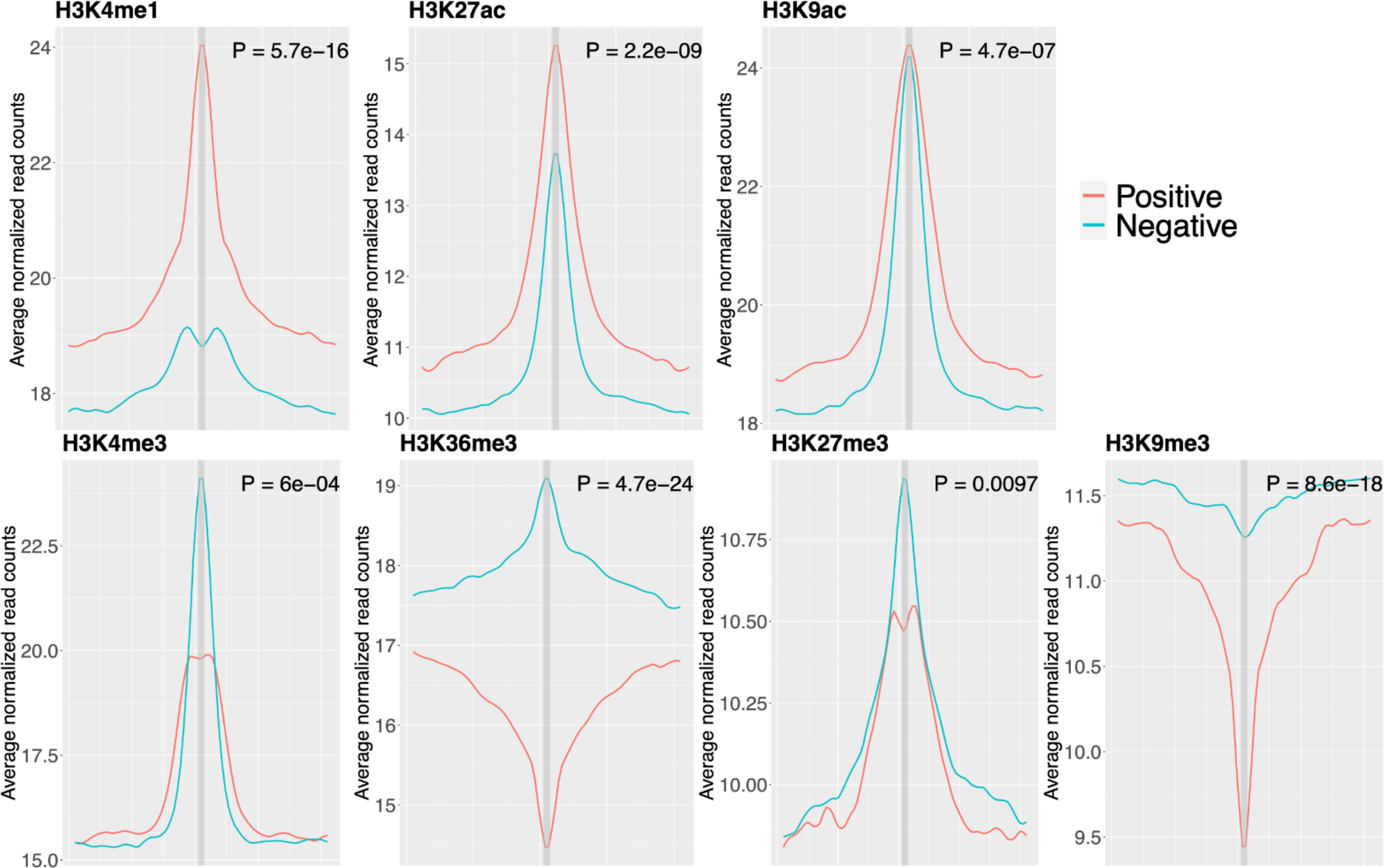
Distribution pattern of histone modification around 5hmC peaks. EB 5hmC peaks is collected from “Forebrain Organoid” 5hmC-seq data and ChIP-seq data in “Brain Angular Gyrus” from seven histone marks are collected from Roadmap Epigenomics. Histone features are obtained and averaged in the neighborhood of all 5hmC peaks for the positive and negative set respectively. Specifically, histone features are created by segmenting an extended genomic region of 10kb both upstream and downstream of each 5hmC peak into 41 1kb windows with a sliding size 500bp and counting reads for each 1kb windows. For each histone mark, Wilcoxon rank-sum test is performed to test the distribution difference of histone features between positive and negative 5hmC peaks and pvalue is reported.

Moreover, the enrichment patterns of the seven histone marks exhibit variations (**Fig. 2**). Active enhancer marks such as H3K4me1, H3K27ac and H3K9ac display a consistent pattern, where the histone features are consistently higher in the positive 5hmC peaks than in negative ones. Notably, H3K4me1 shows the most significant differential enrichment of histone features between positive and negative peaks (pvalue = 5.728 × 10^-16^), followed by H3K27ac (pvalue = 2.169 × 10^-9^) and H3K9ac (pvalue = 4.682 × 10^-7^). This observation aligns with the previous findings that 5hmC is significantly enriched in histone marks associated with enhancers, such as H3K4me1 and H3K27ac (Stroud et al., 2011). Interestingly, the enrichment pattern is opposite for H3K4me3 compared to the active enhancer marks. H3K4me3 is more enriched in negative 5hmC peaks than in positive ones (pvalue = 5.982 × 10^-4^). A similar trend and shape are observed for H3K27me3 (pvalue = 9.690 × 10^-3^). Although the histones features are higher in negative 5hmC peaks than in positive peaks for both H3K9me3 (pvalue = 8.553 × 10^-18^) and H3K36me3 (pvalue = 4.708 × 10^-24^), the enrichment patterns differ. H3K9me3 exhibits a valley-shaped pattern for both positive and negative peaks, while H3K36me3 shows a peak-shaped pattern for negative peaks and a valley-shaped pattern for positive peaks. Comparing to H3K27me3, which is considered as a temporary repression signal, H3K9me3, another repressive mark viewed as a permanent repression signal, shows the same direction but different enrichment patterns (Kim & Kim, 2012). Furthermore, the overall enrichment level of active histone marks is higher than that of repressive histone marks. Taking positive peaks as an example, the average 5hmC read counts in the center of the positive peaks are approximately 24 for H3K4me1, 15 for H3K27ac, 24 for H3K9ac, 20 for H3K4me3 and 14 for H3K36me3 compared to 10 for H3K27me3 and 9 for H3K9me3. Additionally, the change of enrichment from distal windows to the center window of active histone marks is more significant than that of repressive histone marks. For positive peaks, the average 5hmC read counts increases by approximately 5 from the distal 20^th^ window to the center window for H3K4me1, 4 for H3K27ac, 5 for H3K9ac, 5 for H3K4me3, while they decrease by approximately 3 for H3K36me3. In contrast, the average 5hmC read counts increases by only 0.5 for H3K27me3 and decreases by 2 for H3K9me3.

### Evaluating the predictive power of histone marks

Motivated by the observation that the enrichment of histone marks shows differential distribution between positive and negative 5hmC peaks, we further explore the predictive power of each histone mark in classifying positive and negative 5hmC peaks. Identifying the most influential histone marks is crucial for reducing the model complexity especially when dealing with multiple histone marks in the matched tissues or cell types. Specifically, we choose the EB 5hmC peaks from “Forebrain Organoid” and ChIP-seq data from all brain regions of seven histone marks in Roadmap Epigenomics (**Supplement Table S1**) and evaluate the predictive performance for each histone mark in terms of AUROC and AUPRC. Consequently, we find that H3K4me1 and H3K4me3 show higher AUROC than other histone marks (**Supplementary Fig. S1**). In addition, the two histone marks are most prevalent across multiple tissue and cell types in consortiums such as ENCODE and Roadmap Epigenomics. As a result, we only include the two histone marks in the subsequent experiments.

### Comparing unimodal and multimodal Deep5hmC

As Deep5hmC is a multimodal model comprising both sequence and histone modalities, we conduct an ablation study to demonstrate that incorporating the histone modality leads to improved prediction performance of 5hmC modification. For this purpose, we compare two unimodal models of Deep5hmC: Deep5hmC-Seq and Deep5hmC-His to the default multimodal Deep5hmC. Without loss of generality, we utilize the same EB 5hmC peaks from “Forebrain Organoid” and ChIP-seq data from two histone marks: H3K4me1 and H3K4me3 collected from all brain regions in Roadmap Epigenomics (**Supplementary Table S2**). The results indicate that Deep5hmC achieves the best performance with an AUROC of 0.92, followed by Deep5hmC-Seq with an AUROC of 0.89. Deep5hmC-His lags with an AUROC of 0.68 (**Fig. 3A**). The observation indicates that integrating both sequence and histone modalities indeed enhances the prediction performance, although using histone modality alone achieve only moderate predictive ability. Similar trends are observed when measuring prediction performance by AUPRC (**Fig. 3B**). It is evident that multimodal Deep5hmC unequivocally outperforms unimodal Deep5hmC, whether Deep5hmC-Seq or Deep5hmC-His, in terms of both AUROC and AUPRC. Given the superior performance of multimodal Deep5hmC, we will use it as the default implementation in subsequent sections.

**Figure 3.**
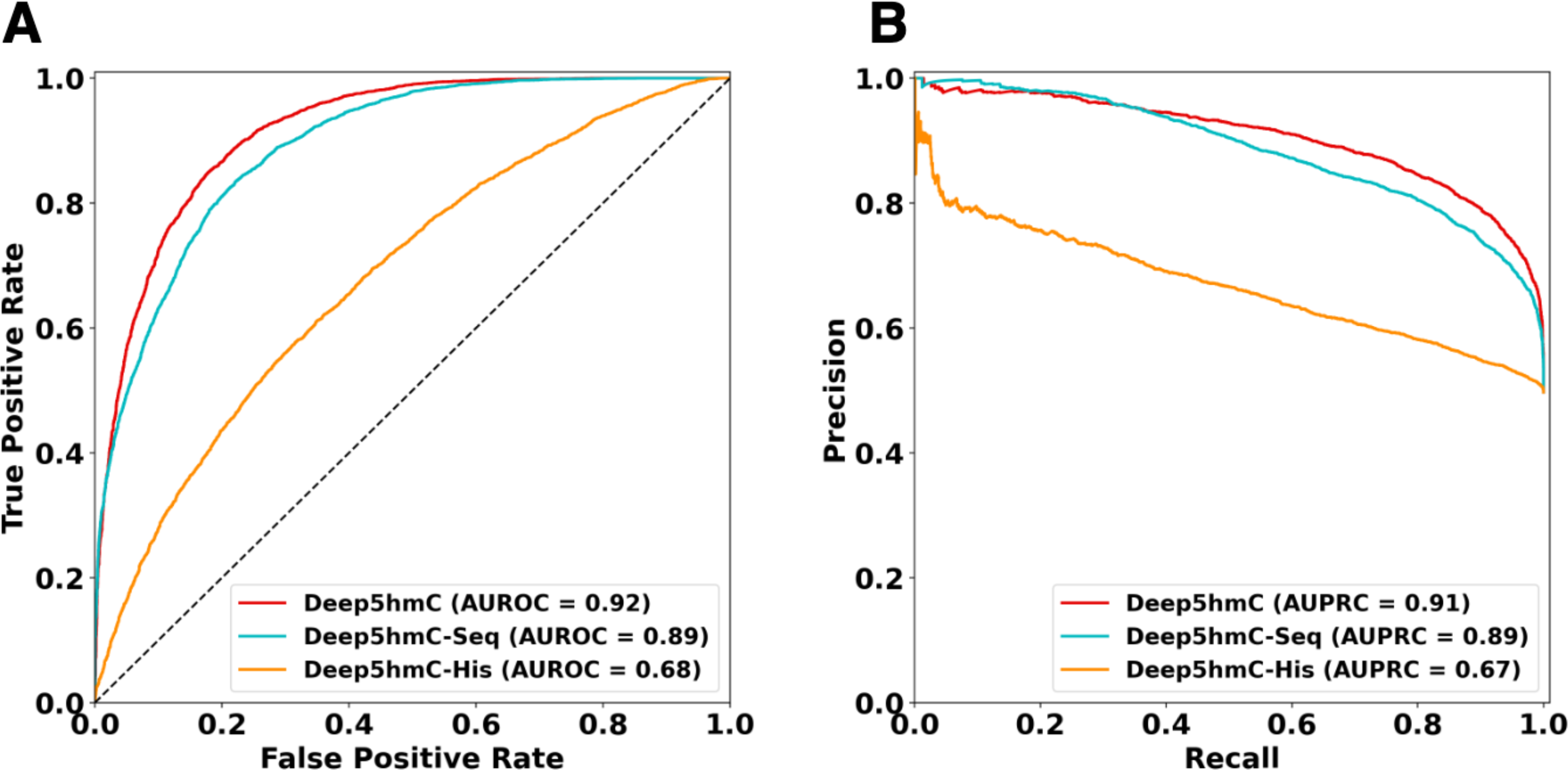
Comparison of unimodal and multimodal Deep5hmC in predicting of 5hmC modification sites. Two unimodal models of Deep5hmC: Deep5hmC-Seq using only DNA sequence as the model input and Deep5hmC-His using only histone features as the model input are compared to the default multimodal Deep5hmC using both DNA sequence and histone features as the model input. 5hmC peaks from “Forebrain Organoid” and two histone marks: H3K4me1 and H3K4me3 ChIP-seq data in all brain regions from Roadmap Epigenomics are used as the training set. **A.** AUROC reported for three compared methods. **B.** AUPRC reported for three compared methods.

### Evaluating Deep5hmC for predicting binary 5hmC modification sites

To demonstrate the superiority of Deep5hmC to existing methods, we further conduct a comparison between the binary version of Deep5hmC, named Deep5hmC-binary, and competing models, which include DeepSEA-Transfer, DeepSEA-Retrain, and Random Forest to predict the binary 5hmC modification sites (i.e. normalized peaks), using the same training, validation, and testing sets as outlined in the “cross-chromosomal” strategy (More details in the Methods section) for both “Forebrain Organoid” and “Human Tissues”. We adopt DeepSEA as the representative of deep learning approaches for its renowned use of genomic sequence to predict epigenetic signals, which has demonstrated superior performance (Zhou & Troyanskaya, 2015). Given that DeepSEA is a multi-task model predicting epigenetic signals across various tissues and cell types simultaneously, we customize it into single-task model in the output layer for a fair comparison to Deep5hmC. In addition, we design two versions of DeepSEA: DeepSEA-Transfer and DeepSEA-Retrain, both sharing the same network architectures as DeepSEA but differing in the training process. Specifically, DeepSEA-Transfer is a transfer learning model built upon the pretrained DeepSEA, with fine-tuning applied exclusively to the last fully connected layer. In contrast, DeepSEA-Retrain starts the model training from the scratch, updating all model parameters. To represent conventional machine learning models, we select Random Forest for its robust performance. Following prior work, we adopt 3-mer frequency of genomic sequence, which result in 64 features for Random Forest (Ahmed et al., 2020).

For “Forebrain Organoid”, we present the AUROC and AUPRC of all models across four developmental time points: day 8 embryoid bodies (EB), day 56 (D56), day 84 (D84), day 112 (D112) of healthy forebrain organoid (**Fig. 4**). Consequently, Deep5hmC-binary consistently obtains the highest AUROC (0.94 for EB; 0.95 for D56; 0.96 for D84; 0.97 for D112) among all methods and developmental stages (**Fig. 4A**). Following closely is DeepSEA-Retrain (0.90 for EB; 0.92 for D56; 0.92 for D84; 0.92 for D112). The observation indicates that enhanced prediction performance can be achieved by leveraging tissue-matched histone modification through a comparison between Deep5hmC-binary and DeepSEA-Retrain, the latter being a CNN model solely relying on genomic sequence as the input. DeepSEA-Retrain outperforms Random Forest (AUROC = 0.85 for EB; 0.88 for D56; 0.89 for D84; 0.90 for D112) and its counterpart DeepSEA-Transfer, which records the lowest overall AUROC (AUROC = 0.86 for EB; 0.85 for D56; 0.83 for D84; 0.83 for D112). The superiority of DeepSEA-Retrain over Random Forest underscores the advantage of deep learning model in capturing nonlinear and high-order dependencies in the genomic sequence compared to k-mer frequency. The observation also suggests that DeepSEA benefits more from retraining the model than relying on the pretrained model. The initial pretraining of DeepSEA on hundreds of tissue/cell type-specific factors may not be optimal for predicting 5hmC modification, which is another epigenetic factor, emphasizing the importance of context matching, where training and testing data belong to the same domain. Moreover, consistent trends are observed across all methods when evaluated using AUPRC. (**Fig. 4B**).

**Figure 4.**
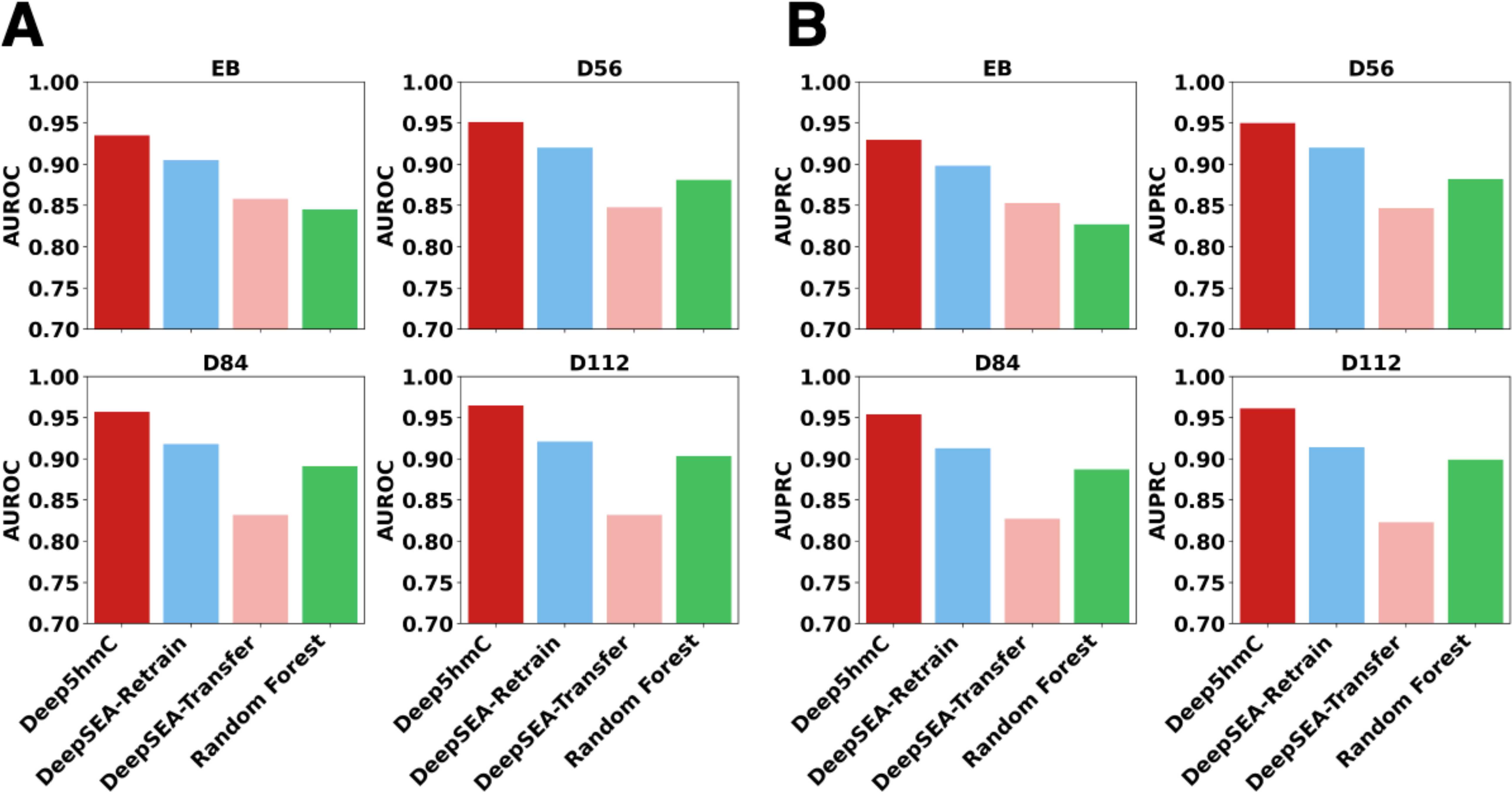
Evaluating Deep5hmC-binary for predicting binary 5hmC modification sites using “Forebrain Organoid” data. **A.** AUROC are reported for all compared methods across 4 developmental stages in “Forebrain Organoid”. **B.** AUPRC are reported for all compared methods across 4 developmental stages in “Forebrain Organoid”.

For “Human Tissues”, we present that both AUROC and AUPRC of all compared methods across 17 human tissues (**Fig. 5**). The evaluation of two methods is extended through a Wilcoxon rank-sum test on AUROC/AUPRC across 17 human tissues, aiming to determine the statistical significance of differences in prediction performance. Consequently, Deep5hmC-binary emerges as the top-performed method, followed by DeepSEA-Retrain, while DeepSEA-Transfer and Random Forest exhibit comparable performance (median AUROC=0.96 for Deep5hmC-binary; 0.90 for DeepSEA-Retrain; 0.84 for DeepSEA-Transfer; 0.85 for Random Forest) (**Fig. 5A**). The superiority of Deep5hmC over other methods is also statistically significant (p-value = 1.548 × 10^-6^ for Deep5hmC-binary versus DeepSEA-Retrain; 6.455 × 10^-7^ for Deep5hmC-binary versus DeepSEA-Transfer; 6.455X 10^-7^ for Deep5hmC versus Random Forest). Consistent with the findings in “Forebrain Organoid”, DeepSEA-Retrain significantly outperforms DeepSEA-Transfer (pvalue = 3.923 × 10^-6^) and DeepSEA-Transfer holds comparable performance to Random Forest. Moreover, the trend is maintained when assessing prediction performance using AUPRC (**Fig. 5B**). Conclusively, the comprehensive evalaution focusing on tissue-specific predictions underscores that Deep5hmC-binary possesses a distinct advantage over existing methods concerning binary 5hmC modification sites.

**Figure 5.**
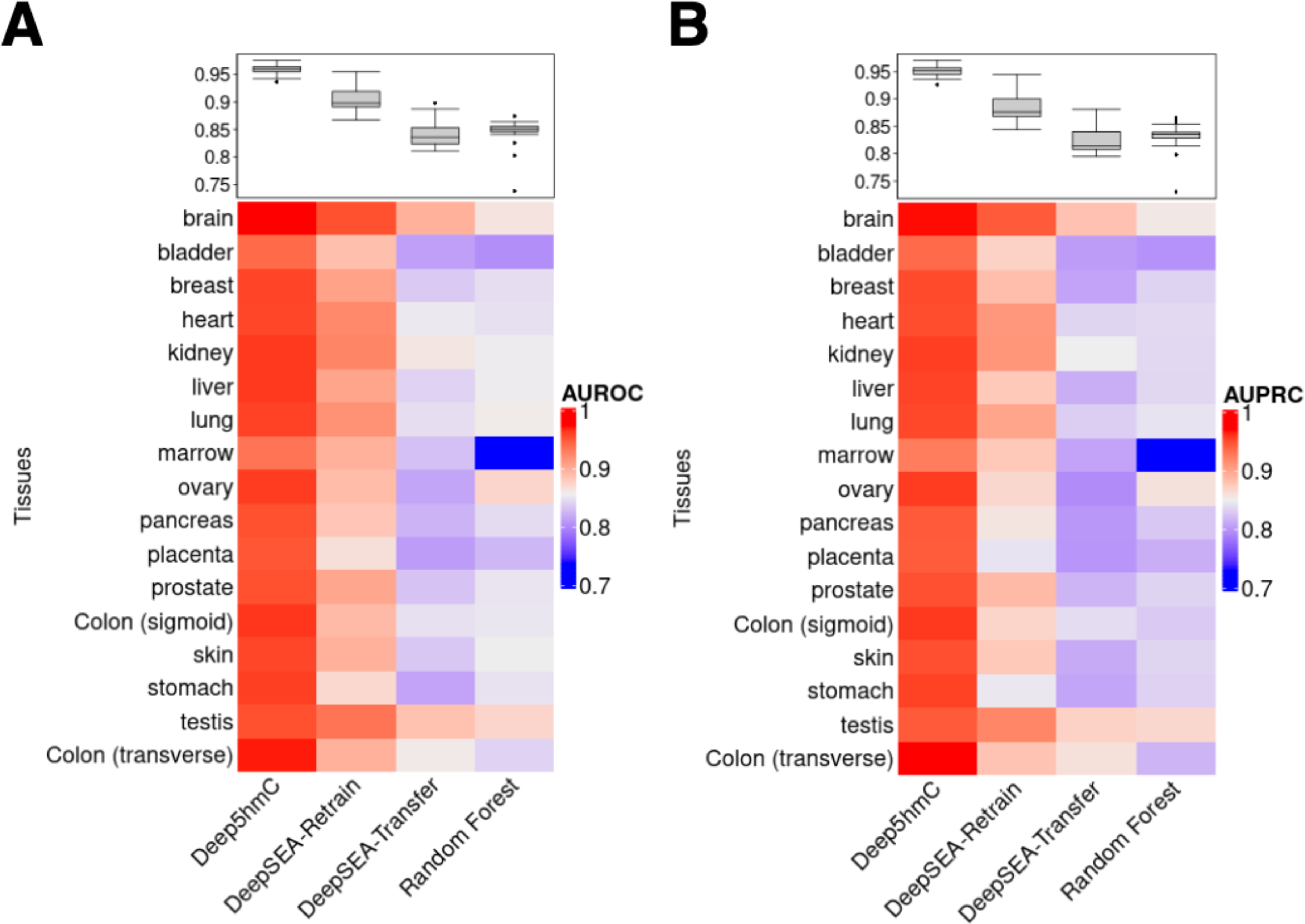
Evaluating Deep5hmC-binary for predicting binary 5hmC modification sites using “Human Tissues” data. **A.** AUROC are reported for all compared methods across 17 human tissues in “Human Tissues”. **B.** AURPC are reported for all compared methods across 17 human tissues in “Human Tissues”.

### Evaluating Deep5hmC for predicting continuous 5hmC modification

Using the same two datasets as aforementioned in predicting binary 5hmC modification, we extend the evaluation to the continuous version of Deep5hmC, named Deep5hmC-cont, along with other competing methods for predicting continuous 5hmC modification, quantified by normalized 5hmC read counts. To mitigate the impact of outliers and normalize the count data towards a normal distribution, we perform a log-transformation on the 5hmC read counts. Evaluation of prediction performance is conducted using Pearson correlation coefficient (R) and mean square error (MSE), which are calculated between the observed 5hmC read counts and predicted ones on the logarithm scale.

For “Forebrain Organoid”, Deep5hmC-cont demonstrates the highest R across all developmental stages (R=0.870 for EB; 0.876 for D56; 0.893 for D84; 0.876 for D112) (**Fig. 6A**). Following closely is DeepSEA-Retrain, showing comparable performance with Random Forest (Rs=0.842, 0.843, 0.857 and 0.790 for DeepSEA-Retrain; 0.829, 0.825, 0.830 and 0.828 for Random Forest). Notably, DeepSEA-Transfer remains potent, yet it exhibits the least favorable performance with Rs of 0.756, 0.693, 0.752 and 0.653 across four time points. In addition, Deep5hmC-cont obtains the lowest MSE in 3 out of 4 time points (**Supplementary Fig. S2A**).

**Figure 6.**
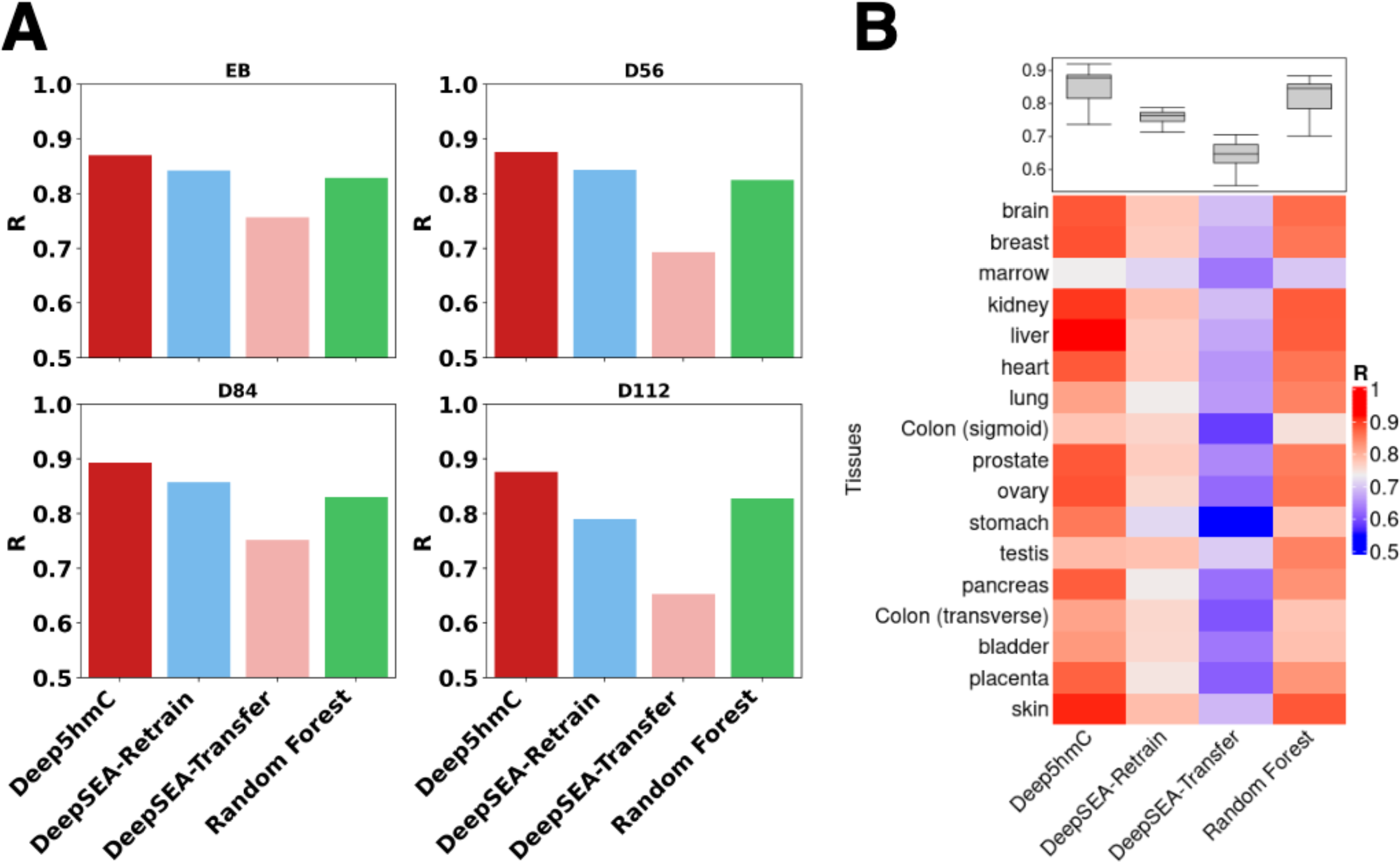
Evaluating Deep5hmC-cont for predicting continuous 5hmC modification. **A.** Pearson correlation coefficients (Rs) are reported for all compared methods across 4 developmental stages in “Forebrain Organoid”. **B.** Pearson correlation coefficients (Rs) are reported for all compared methods across 17 human tissues in “Human Tissues”.

For “Human Tissues”, Deep5hmC-cont excels by emerging as the top-performed method in 15 out of 17 tissues in terms of Rs and achieving the highest median of R across 17 tissues (median R=0.88 for Deep5hmC-cont; 0.76 for DeepSEA-Retrain; 0.65 for DeepSEA-Transfer; R=0.85 for Random Forest). (**Fig. 6B**). Random Forest rank second, followed by DeepSEA-Retrain, while DeepSEA-Transfer exhibits the least favorable performance. Upon comparing the Deep5hmC-cont to other methods using Wilcoxon rank-sum test on the Rs from 17 tissues, we find that Deep5hmC-cont has comparable performance to Random Forest while enjoys a substantial advantage over DeepSEA-Retrain and DeepSEA-Transfer (pvalue = 1.318 × 10^-5^ for Deep5hmC-cont versus DeepSEA-Retrain; 6.455X 10^-7^ for Deep5hmC-cont versus DeepSEA-Transfer; 0.079 for Deep5hmC-cont versus Random Forest). Once again, DeepSEA-Retrain significantly performs better than DeepSEA-Transfer (pvalue = 6.455 × 10^-7^). Additionally, Deep5hmC-cont is comparable to Random Forest and achieves the lowest MSE in 8 out of the 17 tissues, as well as second lowest MSE in 6 out of the 17 tissues (**Supplementary Fig. S2B**). These observations affirm that Deep5hmC-cont accurately predict the tissue-specific continuous 5hmc modification, showcasing its improvement over other methods, especially for “Forebrain Organoid”.

### Evaluating Deep5hmC for predicting gene expression

The positive correlation between 5hmC modification in the gene body and gene expression has been demonstrated in both mouse brain and human tissues (He et al., 2021; Mellén et al., 2012). To quantify the predictive power for gene expression using 5hmC modification, we introduce Deep5hmC-gene, a module within the Deep5hmC framework, designed to predict gene expression by leveraging continuous predicted 5hmC modification from Deep5hmC-cont. Specifically, Deep5hmC-gene employs a three-step approach. First, it segments each gene body into nonadjacent 1kb windows. For gene bodies less than 1kb or with the last window less than 1kb, padding is applied to ensure each window is 1kb. Next, Deep5hmC-gene utilizes the pretrained Deep5hmC-cont to predict the 5hmC counts for each 1kb window. Finally, it aggregates all predicted 5hmC counts within each gene body to generate the predicted gene expression. To evaluate the effectiveness of Deep5hmC-gene in predicting gene expression, we evaluate it on both “Forebrain Organoid” and “Human Tissues”, which provide a comprehensive set of tissue-specific paired 5hmC-seq data and RNA-seq data. Gene expression measured by RNA-seq data, in terms of read counts, serves as the gold standard. The evaluation is based on the pearson Correlation Coefficient (R) calculated between predicted and observed gene expression. In addition, we report Rs calculated between predicted and observed 5hmc read counts in all gene bodies, as the predicted gene expression is quantified by predicted 5hmC read counts in gene bodies.

As a result, Deep5hmC-gene achieves a high R value of 0.95 between the predicted and observed 5hmC read counts in all gene bodies for EB in “Forebrain Organoid” (**Fig. 7A**). Leveraging this accurate prediction of 5hmC in gene bodies, Deep5hmC-gene obtains a substantial R value of 0.54 between predicted and observed gene expression for EB in “Forebrain Organoid” (**Fig. 7B**). Extending the analysis to all four time points in the “Forebrain Organoid” reveals consistent results, with Deep5hmC-gene accurately predicting 5hmC read counts in gene bodies (R within the range of 0.94 to 0.95) (**Fig. 7C**) and gene expression (R within the range of 0.54 to 0.62) (**Fig. 7D**). The MSE calculated between predicted and observed 5hmC read counts in gene bodies, as well as between predicted and observed gene expression for “Forebrain Organoid” can be found in the **Supplementary Fig S3.A,B**.

**Figure 7.**
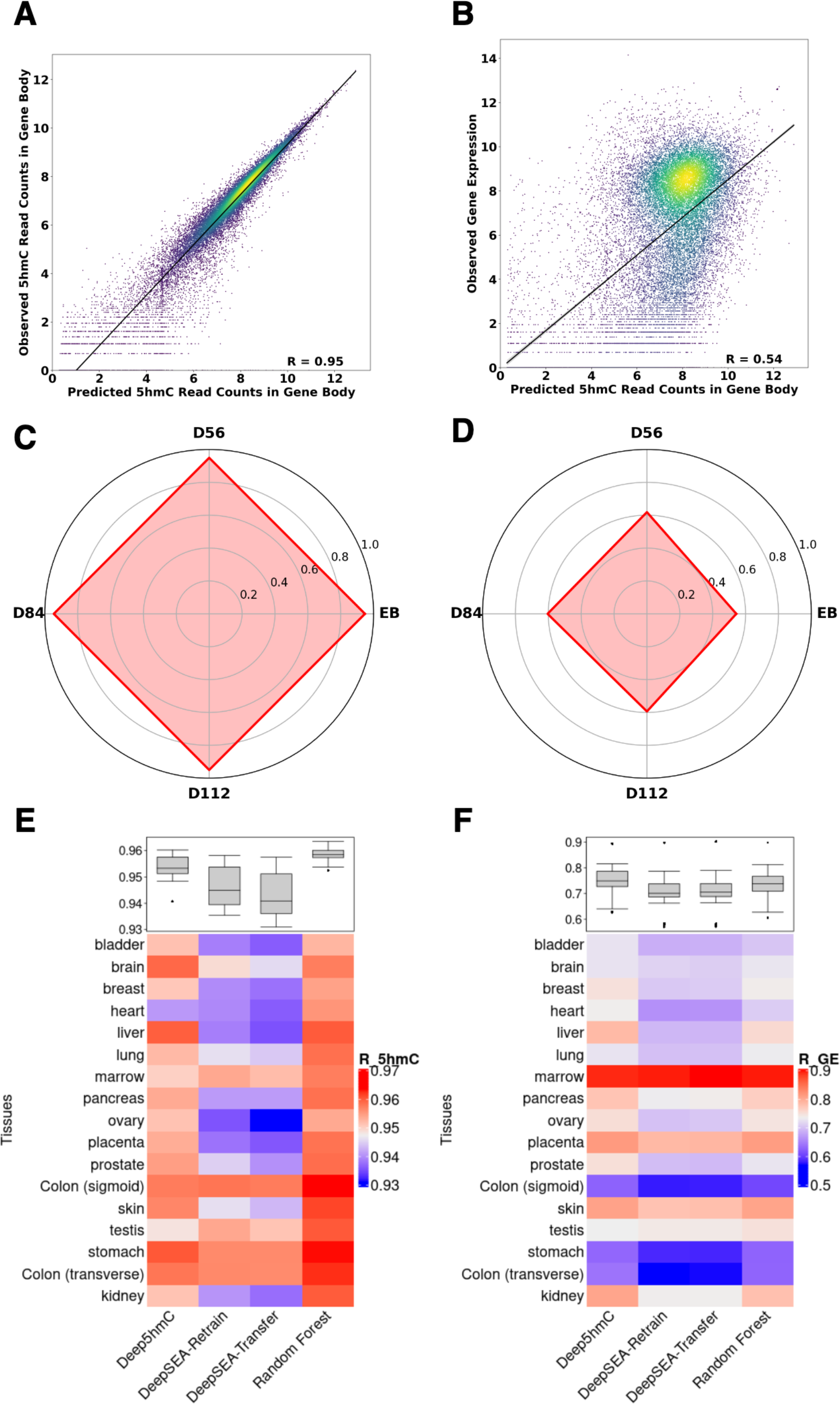
Evaluating Deep5hmC-gene for predicting gene expression. **A.** Correlation between the predicted and observed 5hmC read counts in all gene bodies for EB in “Forebrain Organoid”. **B.** Correlation between predicted and observed gene expression for EB in “Forebrain Organoid”. **C.** Pearson correlation coefficients (Rs) calculated between the predicted and observed 5hmC read counts in all gene bodies for 4 developmental stages in “Forebrain Organoid”. **D.** Pearson correlation coefficients (Rs) calculated between the predicted and observed gene expression for 4 time points in “Forebrain Organoid”. **E.** Pearson correlation coefficients (Rs) calculated between the predicted and observed 5hmC read counts in all gene bodies for 17 human tissues in “Human Tissues”. **F.** Pearson correlation coefficients (Rs) calculated between the predicted and observed gene expression for 17 human tissues in “Human Tissues”.

Moreover, we benchmark Deep5hmC-gene against other competing methods for “Human Tissues”. All approaches demonstrate high prediction accuracy for 5hmC read counts in gene bodies in terms of R (**Fig. 7E**). Deep5hmC-gene exhibits comparable performance to other methods, with a median R value of 0.95 compared to 0.94 for DeepSEA-Retrain, 0.94 for DeepSEA-Transfer and 0.96 for Random Forest. Notably, Deep5hmC-gene exhibits the smallest MSE (**Supplementary Fig S3.C**). The prediction accuracy for gene expression of all methods declines and shows tissue-specific variability (**Fig. 7F**). Deep5hmC-gene performs best, achieving the highest median of R value of 0.75 compared to 0.74 for Random Forest and 0.70 for both DeepSEA-Retrain and DeepSEA-Transfer. Notably, the R values of most tissues falls within the range of 0.7 to 0.8 across all methods, confirming that using 5hmC read counts can accurately predict gene expression. Once again, Deep5hmC-gene achieves the smallest MSE of predicted and observed gene expression for “Human Tissues” (**Supplementary Fig S3.D**). Overall, the exploration demonstrates that leveraging 5hmC read counts in gene bodies facilitates accurate prediction of gene expression, potentially linking DNA methylation and gene expression in a tissue-specific gene regulatory context.

### Evaluating Deep5hmC for predicting differential hydroxymethylated regions

Thus far, we have demonstrated the efficacy of Deep5hmC in predicting tissue-specific 5hmC modification, quantified by both 5hmC peak and continuous 5hmC reads. However, extending our study to a case-control design allows us to explore differential hydroxymethylated regions (DhMRs). In this scenario, DhMRs are regions enriched or present only in one disease/treatment condition, while absent or depleted in the control condition, or vice versa. Identified DhMRs can be potential biomarkers for disease prevention, diagnosis and treatment. We formularize the DhMR identification as a binary classification task, where each genomic region (i.e., 1kb) is labelled and predicted as DhMR or non-DhMR. This module for predicting DhMRs is named “Deep5hmC-diff”. Deep5hmC-diff utilizes Deep5hmC-His modality, masking Deep5hmC-Seq modality, as the genomic sequence is shared between two conditions in the same genomic regions. The training set for Deep5hmC-diff is created by performing differential peak analysis using tools such as DESeq2 (Love et al., 2014) and ChIPComp (Chen et al., 2015) on unioned 5hmC peaks from all samples in both case and control conditions. Subsequently, Deep5hmC-diff is trained and tested using the labeled DhMRs.

To demonstrate the feasibility of Deep5hmC-diff, we focus on an Alzheimer’s disease study, specifically “Kentucky AD”, which profiles 5hmC-seq for 3 AD and 3 healthy controls. We employ DESeq2 (Love et al., 2014) to identify DhMRs (e.g., FDR<0.1) and non-DhMRs (e.g., FDR>0.5), resulting in 4330 differential DhMRs and 4025 non-DhMRs. The histone features are derived from H3K27ac and H3K4me3 ChIP-seq data from Rush Alzheimer’s Disease Study, available on ENCODE portal (Sloan et al., 2016). H3K27ac is used as an alternative of H3K4me1 given the unavailability of H3K4me1 data, and both H3K27ac and H3K4me1 are active enhancer marks. For each histone mark, we select one ChIP-seq data with matched gender, age for the 5hmC-seq in AD group, diagnosed with “Alzheimer’s disease” and another ChIP-seq data with matched gender, age for the 5hmC-seq in healthy control group, diagnosed with “No Cognitive Impairment”.

We employ the same “cross-chromosomal” strategy to create training, validation and testing sets. Deep5hmC-diff achieves an AUROC of 0.67 (**Fig. 8A**) and AUPRC of 0.73 (**Fig. 8B**), demonstrating predictive power for identifying DhMRs by leveraging histone modification data. There is potential for further improvement by incorporating additional histone marks or other epigenetic factors such as chromatin accessibility and transcription factor binding. In addition to evaluating Deep5hmC-diff using the “cross-chromosomal” strategy, we extend its application by conducting a genome-wide screening for *de nove* DhMRs, which may not be present in the 5hmC-seq data potentially due to lacking sufficient sequencing depths or technical bias etc. For this purpose, we utilize all labelled DhMRs and non-DhMRs peaks from “Kentucky AD” to train the Deep5hmC-diff model. The entire human genome is then segmented into non-overlapping 1kb windows, serving as the testing set. Each 1kb window, considered a candidate genomic region, is assigned a predictive probability of being a DhMR or not, using a cutoff at 0.5. The distributions of *de novo* DhMRs is found to be consistent with those from ‘Kentucky AD’ across different genomic features, including Introns, Intergenic Regions, Promoters, Exons, immediate Downstream, 5UTRs and 3UTRs (**Supplement Fig S4A, B**). Of particular interest is the evaluation of whether Deep5hmC-diff can identify *de novo* DhMRs within key functional genomic sites associated with AD. We focus on three causal genes associated with early on-set AD, which include *PSEN1* (chr14:73603143–73690399), *PSEN2* (chr1:227058273-227083804) and *APP* (chr21:27252861-27543138) as well as one causal gene *APOE* (chr19:45409039-45412650) associated with late on-set AD. The predicted probability within the gene bodies, and upstream and downstream 5kb of the gene bodies are plotted (**Fig. 8C**). Deep5hmC-diff successfully identifies multiple *de novo* DhMRs with more DhMR found for *APP* and *PSEN2* than *APOE* and *PSEN1*. As 5hmC modification is positively correlated with gene expression, we conduct differential expression analysis to validate the identified differential *de novo* DhMRs. We collect matched RNA-seq data for AD and healthy controls from “Kentucky AD” and perform differential expression analysis using DESeq2 (Love, Huber, & Anders, 2014). Consequently, three out of the four causal genes show differential expression (FDR = 0.019 for *APP*; 0.031 for *PSEN1*; 0.028 for *PSEN2*), supporting the findings of predicted DhMRs in the three genes. These observations indicate that Deep5hmC-diff can be a valuable tool for identifying novel DhMRs in a case-control study.

**Figure 8.**
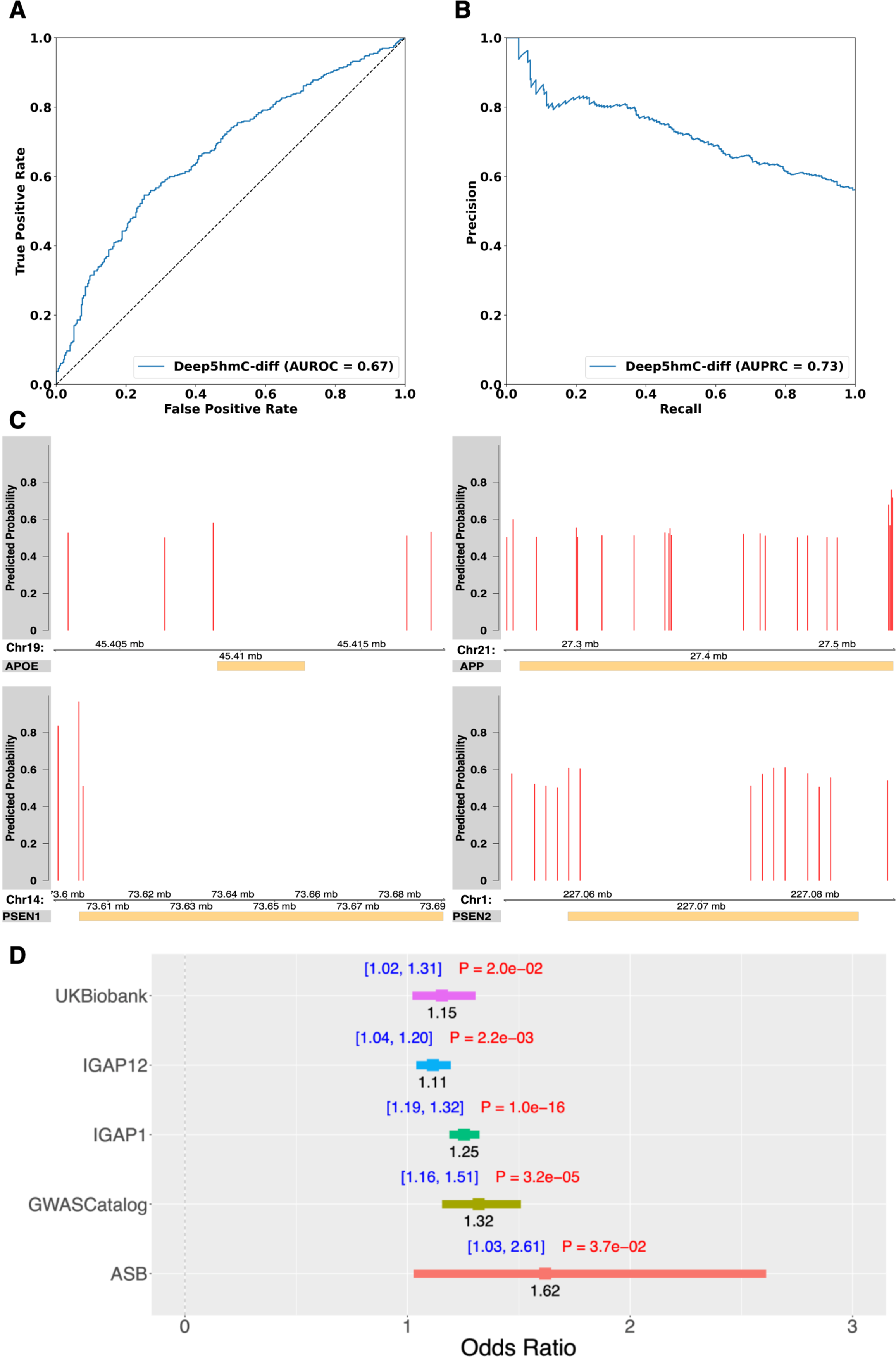
Applying Deep5hmC-diff in a case-control study of Alzheimer’s disease. **A.** AUROC is reported for predicting differential hydroxymethylated regions (DhMRs) between 3 AD patients and 3 healthy controls in “Kentucky AD”. **B.** AUPRC is reported for predicting DhMRs between AD patients and healthy controls. **C.** Distribution of identified *de novo* DhMRs in three causal genes associated with early on-set AD, which include *PSEN1* (chr14:73603143–73690399), *PSEN2* (chr1:227058273-227083804) and *APP* (chr21:27252861-27543138) as well as one causal gene *APOE* (chr19:45409039-45412650) associated with late on-set AD. **D.** SNP enrichment analysis to evaluate the enrichment of AD-associated SNPs in *de novo* DhMRs. Positive SNPs are collected from five sources including UK Biobank. Association Results Browser (ARB), GWASCatalog, International Genomics of Alzheimer’s Project (IGAP) stage1 and combined stage1 & 2.

The SNP enrichment analysis is designed to assess the enrichment of diseases/traits-associated GWAS SNPs or eQTLs within tissue/cell type-specific epigenetic regions. The analysis is crucial for identifying disease/trait-associated cell types, providing functional annotation and elucidating the role of GWAS SNPs or eQTLs (Agarwal et al., 2023; Chen, Jin, & Qin, 2016; Chen & Qin, 2017; Chen et al., 2019; Kundaje et al., 2015; Wang & Chen, 2022). Recent AD studies have extensively employed SNP enrichment analysis to assess the enrichment of AD-associated SNPs in DhMRs, which helps unravel the functional implications of these AD-associated SNPs in AD pathogenesis (Bernstein et al., 2016). Building on these insights, we conduct SNP enrichment analysis to evaluate the enrichment of AD-associated SNPs in *de novo* DhMRs, which are defined as genome-wide 1kb candidate regions with a predictive probability greater than 0.5 contrasting with non-DhMRs.

Statistically significant AD-associated SNPs, considered positive SNPs, are gathered from five resources, containing summary statistics from GWAS conducted in AD. The first set of positive SNPs is derived from a study named “genome-wide association study by proxy (GWAX)”, comprising 1302 significant SNPs from UK Biobank (pvalue<1×10^-4^) (Liu, Erlich, & Pickrell, 2017). The second set is obtained from GWASCatalog (https://www.ebi.ac.uk/gwas/), including 1108 significant SNPs (p-value<1x 10^-4^). The third set is sourced from the Association Results Browser (ARB) (https://www.ncbi.nlm.nih.gov/projects/gapplus/sgap_plus.htm), containing 111 significant SNPs (p-value<1x 10^-4^). The other two sets of positive SNPs are acquired from International Genomics of Alzheimer’s Project (IGAP) stage1 and combined stage1 & 2, harboring 6225 and 3687 significant SNPs respectively (p-value<1×10^-4^) (Lambert et al., 2013). Moreover, only SNPs in the noncoding regions are considered given the majority of GWAS SNPs are located within noncoding regions. To establish a reliable control group, we generate negative control SNPs at a ratio of 10:1 compared to the positive set, following the strategy from previous work (Chen, Jin, & Qin, 2016; Chen & Qin, 2017). Subsequently, for each variant set, we tally the number of positive/negative SNPs within DhMRs/non-DhMRs and construct a 2 by 2 contingency table. Fisher’s exact test is then employed to calculate the odd ratio (OR), confidence interval (CI) and pvalue for the table. The results reveal that all five sets of positive SNPs exhibit enrichment in the *de novo* DhMRs (OR>1 and pvalue<0.05), suggesting a crucial role of AD-associated SNPs in the pathogenesis of AD through their enrichment in DhMRs.

## Conclusion and Discussion

In this study, we present a comprehensive deep learning framework named Deep5hmC, designed to predict genome-wide landscape of 5-Hydroxymethylcytosine (5hmC). Deep5hmC comprises four distinct modules tailored to specific prediction tasks: Deep5hmC-binary for predicting binary 5hmC peaks; Deep5hmC-cont for predicting continuous 5hmC modification; Deep5hmC-gene for predicting gene expression; Deep5hmC-diff for predicting differential hydroxymethylated regions (DhMRs). Notably, Deep5hmC stands out as a multi-modal deep learning model, which incorporates both DNA genomic sequence and histone modification data to enhance the accuracy of prediction for genome-wide qualitative and quantitative 5hmC modification. The decision to include histone modality stems from a thorough exploration of real data, involving tissue-matched histone ChIP-seq data from seven histone marks in a specific one brain region and one 5hmC-seq data profiled in embryoid body (EB) from forebrain organoid. This exploration reveals distinct distribution patterns between 5hmC peaks and non-peak genomic regions in histone modifications of both active and repressive histone marks, suggesting the informative nature of histone modification features in predicting 5hmC modification. Notably, H3K4me1 and H3K4me3 are identified as the most informative histone marks. To accommodate the histone modality, Deep5hmC employs *n* Convolutional Neural Networks (CNNs), each corresponding to a different histone mark. The output from the histone modality is then integrated with the output from the sequence modality through the MFB fusion layer, resulting in a joint embedding for subsequent predictions. Using an illustrative example with one 5hmC-seq data profiled in EB from a brain organoid, we demonstrate that the multi-modal Deep5hmC outperforms both single-modal Deep5hmC-Seq, utilizing only DNA sequence, and Deep5hmC-His, relying solely on histone modification.

We further employ the multi-modal version of Deep5hmC as the default model for comparative analysis against existing methods, which include Random Forest and two variants of DeepSEA involving fine-tuning or retraining on two comprehensive datasets. These datasets encompass a broad collection of 5hmC-seq across human tissues. One dataset, named “Forebrain Organoid”, comprises matched 5hmC-seq and RNA-seq from four stages during fetal brain development. The other dataset, named “Human Tissues”, includes matched 5hmC-seq and RNA-seq from 17 diverse human tissues. Through an evaluation using the “cross-chromosomal” strategy, Deep5hmC-binary emerges as superior to existing methods, achieving the highest AUROC and AUPRC for predicting binary 5hmC modification sites. Similarly, Deep5hmC-cont attains the highest Pearson correlation coefficient and lowest MSE for predicting continuous 5hmC modification. Moreover, leveraging the predictions from pretrained Deep5hmC-cont, Deep5hmC-gene aggregates all predicted 5hmC counts within the gene body, accurately predicting the gene expression for both “Forebrain Organoid” and “Human Tissues”. This observation underscores the regulatory connection between DNA hydroxymethylation and gene expression in a tissue-specific context.

In addition to predicting 5hmC modification in a single healthy tissue, Deep5hmC-diff enables the prediction of differential hydroxymethylated regions (DhMRs) in a case-control design. where the regions, enriched or present only in one disease/treatment condition but depleted or absent in the control condition (or vice versa), are of particular interest. Demonstrating the feasibility, Deep5hmC-diff is applied to “Kentucky AD” study with matched 5hmC-seq and RNA-seq data for both AD patients and healthy controls. The results not only showcase the accurate prediction of DhMRs using the “cross-chromosomal” strategy but also successfully identify genome-wide *de novo* DhMRs. Notably, multiple *de novo* DhMRs are found in AD causal genes such as *APP*, *APOE*, *PSEN1* and *PSEN2*. These findings are further supported by differential expression analysis using the matched RNA-seq data. In addition, significant SNPs reported to be associated with AD from various studies are found to be enriched in DhMRs, indicating a potential role of DhMRs in AD pathogenesis. Overall, these discoveries underscore the potency and broad applications of Deep5hmC in 5hmC-seq analysis.

Several promising extensions of current work are envisioned. First, the 5hmC-seq data used to train Deep5hmC lacks single-base resolution. The incorporation of high-resolution 5hmC data from advanced technologies such as Tet-assisted bisulfite sequencing (TAB-Seq) and Oxidative bisulfite sequencing (oxBS-Seq), provides an opportunity to extend Deep5hmC’s capability to predict 5hmC modification at the single-base level. To achieve this, we intend to adapt and develop large language models, accommodating the significantly increased training sample size and addressing spatial correlation among single-base modification sites. Secondly, while we have incorporated histone modification as one additional modality for Deep5hmC, other epigenetic factors such as transcription factor binding and chromatin accessibility can be further integrated into the multi-modal deep learning framework. This expansion aims to enhance prediction performance by considering a more comprehensive set of epigenetic features. Furthermore, in our future work, we plan to develop an explainable version of Deep5hmC utilizing attention mechanisms. This will enable the identification of functional interactions between 5hmC and other epigenetic marks, shedding light on their interplay in the regulation of gene expression. This approach seeks to provide a more interpretable and nuanced understanding of the complex relationships within the epigenetic landscape.

## Methods

### Data description and processing

The first dataset, termed “Forebrain Organoid”, includes paired 5hmC-seq data and RNA-seq data across embryoid body (8 days EB) and forebrain organoids cultured over three distinct time points: 56 days (D56), 84 days (D84), and 112 days (D112), designed to model the early development of the fetal brain (Kuehner et al., 2021). The called 5hmC peaks using MACS2 are retrieved from the original publication with GEO accession number GSE151818 (Kuehner et al., 2021). Each 5hmC peak is subsequently standardized into a 1kb window by extending the center of the peak upstream and downstream 500bp. For the acquisition of raw read counts associated with each peak, the raw 5hmC-seq data is downloaded, and bowtie2 (Langmead et al., 2009) is employed to map the reads onto hg19 reference genome. Using R/Bioconductor package “GenomicRanges”, read counts for each 5hmC peak are obtained by overlapping the genomic positions of reads and peaks. Similarly, raw RNA-seq data is obtained, and STAR (Dobin & Gingeras, 2015) is utilized to map the reads onto hg19 reference transcriptome. Read counts for each gene are calculated based on the positional overlap between reads and genes, using R/Bioconductor package “GenomicRanges” and “Rsamtools”. Subsequently, the read counts of biological replicates are averaged after adjusting the sequencing depth.

The second dataset, referred to as “Human Tissues”, comprises paired 5hmC-seq data and RNA-seq data spanning 19 human tissues derived from ten organ systems. The called 5hmC peaks using MACS2 are obtained from the original publication with GEO accession number GSE144530 (Cui et al., 2020). To enhance reliability, we merge the peaks from biological replicates and retain only those peaks appearing in more than two biological replicates. Subsequently, the merged peaks are further standardized into 1kb windows. Raw 5hmC-seq data is downloaded and processed by mapping reads onto hg19 reference genome using bowtie2. Read counts for each 5hmC peak are then calculated based on the mapped genomic positions. Raw RNA-seq data is downloaded and processed using STAR to map reads onto hg19 reference transcriptome. The read counts for each gene are determined by overlapping genomic positions between reads and genes. The read counts of biological replicates are averaged while adjusting the sequencing depth.

The third dataset, titled “Kentucky AD”, is obtained from one the publication which provides the information of 5hmC-seq data in an Alzheimer’s disease study conducted by University of Kentucky Alzheimer’s Disease Research Center with GEO accession number GSE72782 and RNA-seq With SRA accession number SRA060572 (Bernstein et al., 2016). Raw 5hmC-seq data is collected from three prefrontal cortex samples of post-mortem AD patients and three controls with similar age, no history of neurological illness and no significant neuropathology. After data acquisition, read mapping is executed using bowtie2 on the hg19 reference genome, and peak-calling is conducted for each sample using MACS2. Peak standardization and read counting for each peak are carried out employing the aforementioned approaches.

For histone modification data, our primary focus is on acquiring H3K4me1 and H3K4me3 ChIP-seq data that aligned with the tissue or disease condition associated with the 5hmC-seq data. In the case of “Forebrain Organoid”, we compile aligned bed files of brain-related ChIP-seq data from Roadmap Epigenomics (Kundaje et al., 2015) (https://egg2.wustl.edu/roadmap/data/byFileType/alignments/unconsolidated/) (**Supplementary Table S2**). For “Human Tissues”, we carefully select aligned bed files of ChIP-seq data from Roadmap Epigenomics by ensuring a match between ChIP-seq data and 5hmC-seq data based on tissue type. In cases where tissue-matched ChIP-seq data is unavailable at Roadmap Epigenomics, we retrieve it from ENCODE portal (Sloan et al., 2016) (https://www.encodeproject.org/). Owing to the absence of matched histone ChIP-seq data for “Hypothalamus” and “Lymph Nodes” in both databases, we exclude the two tissues, resulting in a total of 17 tissues in “Human tissues” for the subsequent analysis (**Supplementary Table S3**). For “Kentucky AD”, we gather H3K27ac and H3K4me3 ChIP-seq data from Rush Alzheimer’s Disease Study available on ENCODE portal (**Supplementary Table S4**). H3K27ac is used as an alternative for H3K4me1 due to the unavailability of H3K4me1 ChIP-seq data and both H3K27ac and H3K4me1 serve as active enhancer marks. In addition, H3K27ac has been found correlated with 5hmC (Cui et al., 2020). For AD group, we collect mapped ChIP-seq bam files from one individual with matched gender, age and diagnosed with “Alzheimer’s disease”. For healthy control group, we obtain the mapped ChIP-seq bam files from one individual with matched gender, age and diagnosed with “No Cognitive Impairment”.

### Multimodal features

Two types of features are utilized as input for Deep5hmC, which include DNA sequence within the standardized 5hmC peak (i.e., 1kb) and histone modification in the proximity of 5hmC peak. The DNA sequence in each 1kb window undergoes one-hot encoding, adhering to the rule ‘A’: [1,0,0,0], ‘C’: [0,1,0,0], ‘G’: [0,0,1,0] and ‘T’: [0,0,0,1], which result in a 1000 4 matrix representing the sequence feature. For the histone feature, we extend 10kb both upstream and downstream of each 5hmC peak and calculate normalized read counts from matched tissue-specific histone ChIP-seq data in a 1kb window with a sliding size 500bp, yielding the histone feature with dimensions 1×41. In scenarios where *n* matched ChIP-seq datasets are available, histone features from all datasets are horizontally stacked, resulting in a histone feature with dimensions *n*×41.

### Creating labelled data of training, validation and testing

The qualitative prediction is essentially a binary classification task aimed at distinguishing 5hmC peaks from background genomic regions. Specifically, we label standardized 5hmC peaks (i.e., 1kb) with statistical significance from peak-calling results (FDR<0.05) as positive. To choose peaks in the negative set, we apply a series of selection criteria for genome-wide 1kb genomic regions of hg19 reference genome. Initially, negative peaks are required to be within 10kb distance from the positive ones. Additionally, the density distribution of GC content in the negative peaks must match that of the positive ones. Without loss of generality, we maintain an equal number of positive and negative peaks. As a result, the number of positive peaks ranges from 12,596 to 137,488 with a median of 69,322 among 17 human tissues in “Human Tissue” and from 56,036 to 81,050 with a median of 72,745 for “Forebrain Organoid”. To predict differentially hydroxymethylated regions (DhMRs) in “Kentucky AD”, we start by identifying the 5hmC peaks from all samples in both AD and healthy controls. We merge overlapped peaks, standardize and calculate the read counts for merged 5hmC peaks. Next, we employ DESeq2 (Love et al., 2014) to identify DhMRs. Peaks exhibiting statistical significance are deemed as positive (e.g., FDR<0.1) and non-significant peaks as considered as negative (e.g., FDR>0.5). For the quantitative prediction of 5hmC modification, we treat logarithm of normalized 5hmC reads from both peaks and non-peak genomic regions as the outcome. More details regarding sample size from all datasets can be found in **Supplementary Table S6**.

### Network architecture of Deep5hmC

Deep5hmC is essentially a multimodal deep learning model, which consists of three crucial components in the network architecture, which includes (1) an encoder module based on two CNNs; (2) a feature fusion module based on Multi-modal Factorized Bilinear (MFB) pooling approach (Yu et al., 2017) and (3) a prediction module for either binary classification or continuous prediction (**Fig. 2D**).

The encoder module is composed of two unimodal encoders, each responsible for transforming an individual modality to a high-level feature presentation for further processing by subsequent layers in the model. Specifically, two separate and independent CNNs function as the unimodal encoders for DNA sequence and histone modification respectively. The sequence encoder takes the one-hot encoding DNA sequence as input, consisting of three sequential 1-D convolutional layers sharing the same kernel size of 8 and stride of 1, padding of 0, and dilation of 1. The number of filters vary across these layers: 64, 128 and 256. In addition, a max-pooling layer with a kernel size of 4 and a stride of 4 follows each of the first two convolutional layers. The output of last convolutional layer is flattened and connected to two fully connected layers. On the other hand, the histone encoder takes curated histone features from *n* histone marks as the input, where each histone mark *h* is profiled in the dimensions *n_h_*×41. Here, *n_h_* represents the number of matched tissues/cell types or biological replicates of the histone ChIP-seq data. Consequently, the histone encoder takes multimodal histone features as input, with each modality representing a different histone mark (**Fig. 2D**). Each histone mark has its own CNN to extract the high-level features, comprising three 2-D convolutional layers sharing the same kernel size of 3X 3, stride of 1, padding of 1, and varying number of filters: 32, 64, 128. A max-pooling layer with stride of 2 follows each of the first two convolutional layers. The kernel size for the max-pooling layer depends on the *n_h_*. For *n_h_* equals 1, a 1X2 kernel size is chosen, and otherwise 2X 2. Similarly, the output of last convolutional layer is flattened and is connected to two fully connected layers. Finally, the output from *n* CNN, corresponding to *n* histone marks, are concatenated to form the final output of histone module.

The feature fusion module seamlessly integrates the two feature representations derived from the sequence and histone encoders into a unified representation for subsequent prediction (**Fig. 2D**). Specifically, we employ MFB (Yu et al., 2017), designed to efficiently amalgamate features from diverse modalities. Compared to alternative fusion techniques, MFB excels in capturing intricate interactions among multiple modalities while concurrently reducing computational complexity through factorization. Let ***x***_1_ ∊ ℝ^*m*^ denote the feature representation from sequence modality, ***x***_2_ ∊ ℝ^*n*^ represent feature representation from histone modality and ***z*** ∊ ℝ^*o*^ denote the output after fusion module. Notably, *o* is substantially smaller than both *m* and *n*. MFB aims to identify two low-rank factorized matrices 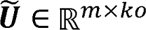 and 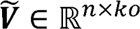, aiming to convert two long vectors of different lengths to two short vectors of same length *ko*, where *k* denotes the latent dimensionality indicating the degree of factorization. The larger *k* is, the more original information can be preserved.

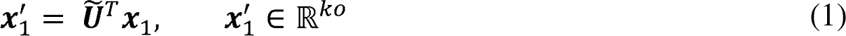

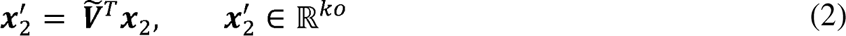

The output of MFB fusion can be represented as follows:

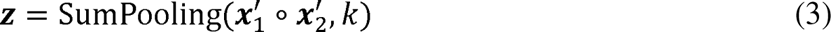

where ° is the element-wise multiplication of two vectors. Following the SumPooling operation, subsequent layers include power normalization (sign(***z***)|***z***|^0.5^) and ℓ_2_ normalization (***z***/|| ***z*** ||) layers. These steps enhance the properties of the fused data, ensuring appropriate scaling and distribution characteristics for subsequent layers.

The prediction module utilizes the output of MFB fusion layer as the input for the two fully connected layers, which are succeeded by the output layer for the prediction. In the output layer, a single node exists for continuous outcome and two nodes are present for binary outcome. In the case of binary outcome (presence or absence of 5hmc peak), the output undergoes a sigmoid function to yield the prediction probability. For continuous outcome (normalized 5hmC read counts), the output directly serves as the prediction. ReLU serves as the activation function across the entire network, excluding the output layer. Additionally, dropout layers with a rate of 0.5 are strategically incorporated to mitigate overfitting.

### Model implementation, training, validation and testing

Deep5hmC is implemented using PyTorch (Paszke et al., 2019) on an NVIDIA A100 GPU system. Utilizing mini-batch gradient descent and the Adam optimizer (Kingma & Ba, 2017), the network is optimized for binary outcome using cross-entropy loss and continuous outcome using mean square error (MSE) respectively. The default learning rate is set to 10^-3^. To improve the efficiency of learning process, warm-up steps and a learning rate decay strategy are incorporated as options. Each model undergoes training for a maximum of 200 epochs, with early stopping implemented if the model performance stagnated over a consecutive 10 epochs. In alignment with the evaluation strategy for DeepSEA, a “cross-chromosomal” strategy is employed to design training, validation and testing sets. Specifically, 5hmc peaks on chromosomes 8 and 9 constitute the testing set, chromosome 7 serves as the validation set, and the remaining chromosomes form the training set.

## Availability of data and materials

The histone ChIP-seq data can be found at Roadmap Epigenomics (https://egg2.wustl.edu/roadmap/data/byFileType/alignments/unconsolidated/) and ENCODE portal (https://www.encodeproject.org/). 5hmC peaks and raw 5hmC-seq data and RNA-seq data of “Forebrain Organoid” can be found from GEO with accession number GSE151818. 5hmC peaks and raw 5hmC-seq data and RNA-seq data of “Human Tissues” can be found from GEO with accession number GSE144530. For “Kentucky AD”, raw 5hmC-seq data can be found from GEO with accession number GSE72782 and RNA-seq with SRA accession number SRA060572.

## Software availability

https://github.com/XinBiostats/Deep5hmC

## Acknowledgements

This work was supported by the following funding sources: NIH grants R35GM142701 to L.C.

## Author contributions

L.C conceived and designed the study. X.M., R.S.T. and L.C. performed the data analysis. X.M. and L.C. wrote the manuscript with input from all the other authors. All authors have read and approved the final manuscript.

## Ethics declarations

The authors declare no competing interests.

## Supplementary information

**Supplementary Figure S1. Evaluating predictive performance of 7 histone marks using ‘Forebrain Organoid’ data. A.** AUROC are reported for all histone marks for EB in “Forebrain organoid”. **B.** AUPRC are reported for all histone marks for EB in “Forebrain Organoid”.

**Supplementary Figure S2. Evaluating Deep5hmC-cont for predicting continuous 5hmC modification. A.** Mean squared error (MSE) are reported for all compared methods across 4 developmental stages in “Forebrain Organoid”. **B.** MSE are reported for all compared methods across 17 human tissues in “Human Tissues”.

**Supplementary Figure S3. Evaluating Deep5hmC-gene for predicting gene expression. A.** Mean squared error (MSE) are calculated between the predicted and observed 5hmC read counts in all gene bodies for 4 developmental stages in “Forebrain Organoid”. **B.** MSE are calculated between the predicted and observed gene expression for 4 developmental stages in “Forebrain Organoid”. **C.** MSE are calculated between the predicted and observed 5hmC read counts in all gene bodies for 17 human tissues in “Human Tissues”. **D.** MSE are calculated between the predicted and observed gene expression for 17 human tissues in “Human Tissues”.

**Supplementary Figure S4. Comparing the distribution of DhMRs in the training set of “Kentucky AD” to genome-wide *de novo* DhMRs across different genomic features. A.** The distribution of DhMRs in the training set of “Kentucky AD” across different genomic features. **B.** The distribution of genome-wide *de novo* DhMRs across different genomic features.

**Supplementary Table S1. Source of ChIP-seq data in “Brain Angular Gyrus” from Roadmap Epigenomics for exploring the distribution pattern of histone modification in the neighborhoods of EB 5hmC peaks from “Forebrain Organoid”.**

**Supplementary Table S2. Source of ChIP-seq data in all brain regions from Roadmap Epigenomics for evaluating the predictive power of seven histone marks and being used by Deep5hmC as histone features (H3K4me1 and H3K4me3) for evaluating 4 developmental stages in “Forebrain Organoid”.**

**Supplementary Table S3. Source of ChIP-seq data from ENCODE being used by Deep5hmC as histone features (H3K4me1 and H3K4me3) for evaluating 17 human tissues in “Human Tissues”.**

**Supplementary Table S4. Source of ChIP-seq data from ENCODE used by Deep5hmC as histone features (H3K27ac and H3K4me3) for predicting DhMRs in “Kentucky AD”.**

**Supplementary Table S5. Summary of sample size for “Brain Organoid”, “Human Tissues” and “Kentucky AD” data.**

