## Supplementary materials for "Deep5hmC: Predicting genome-wide 5-Hydroxymethylcytosine landscape via a multimodal deep learning model"

### Supplementary Figures

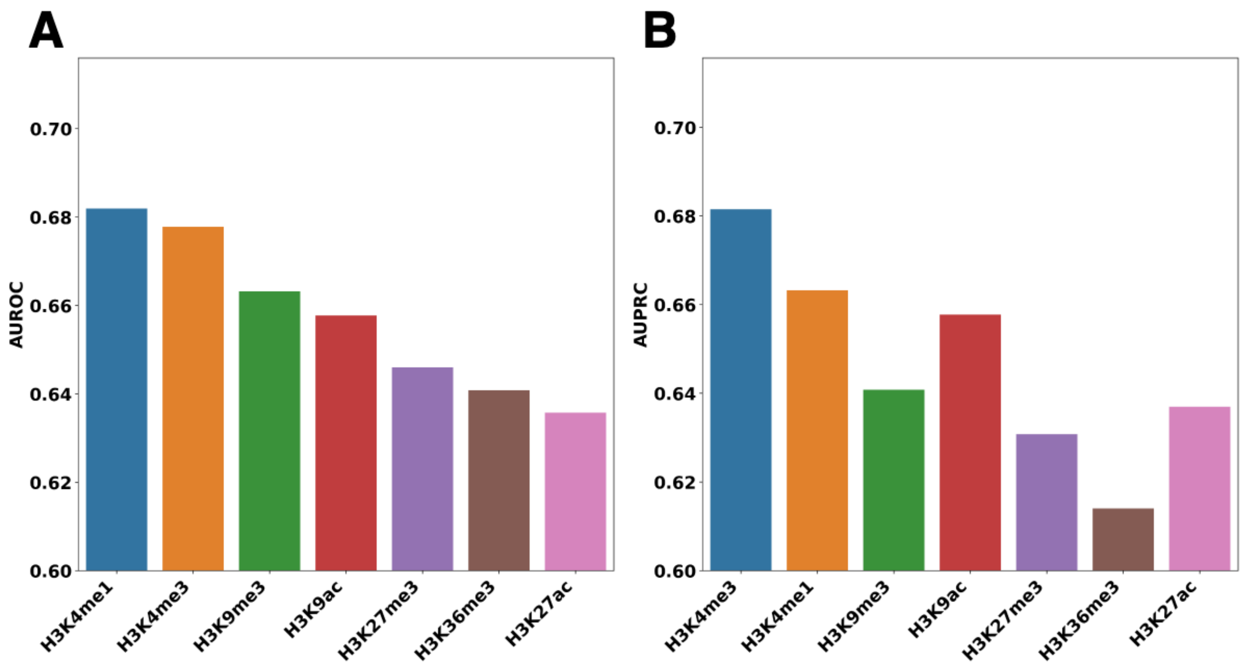


**Supplementary Figure S1. Evaluating predictive performance of 7 histone marks using 'Forebrain organoid' data. A.** AUROC are reported for all histone marks for EB in "Forebrain Organoid". **B.** AUPRC are reported for all histone marks for EB in "Forebrain Organoid".

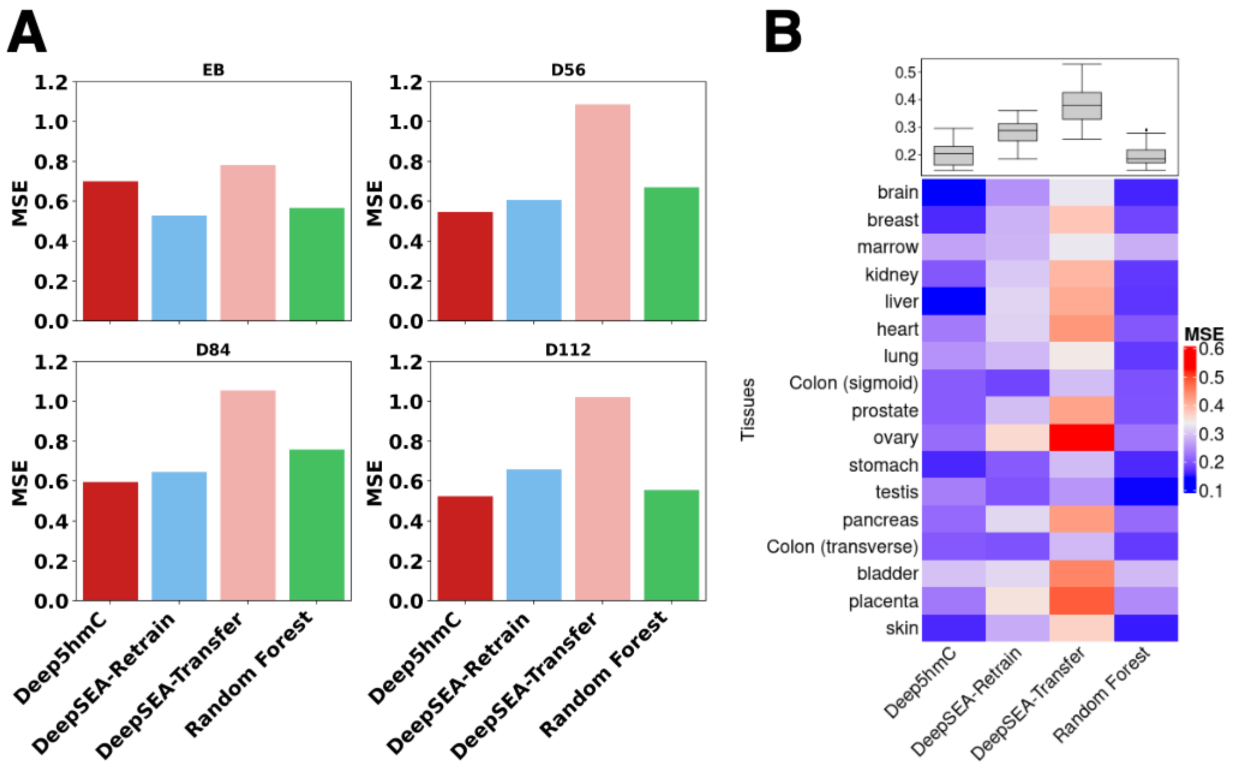


**Supplementary Figure S2. Evaluating Deep5hmC-cont for predicting continuous 5hmC modification.**

**A.** Mean squared error (MSE) are reported for all compared methods across 4 time points in "Forebrain Organoid". **B.** MSE are reported for all compared methods across 17 human tissues in "Human Tissues".

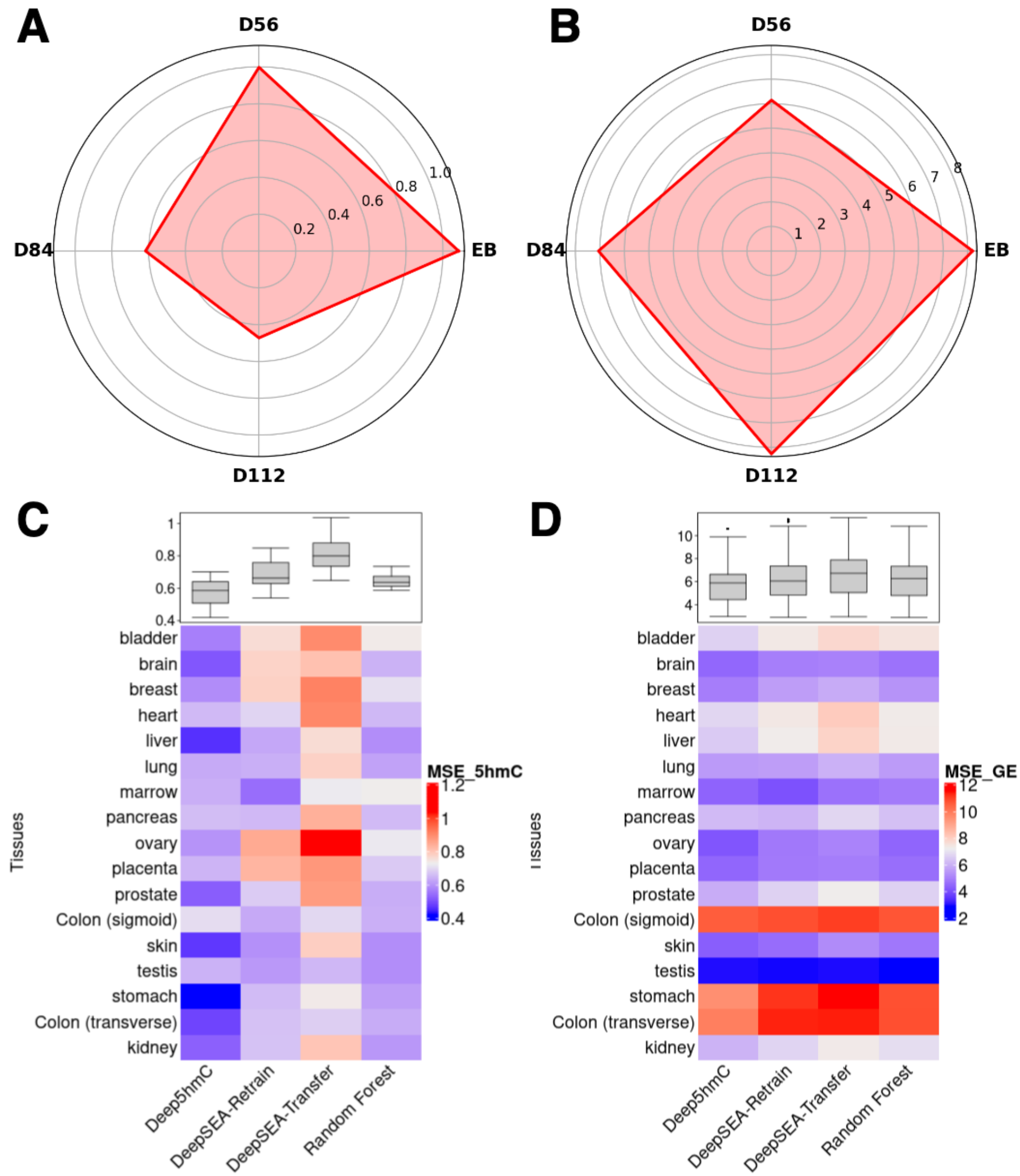


**Supplementary Figure S3. Evaluating Deep5hmC-gene for predicting gene expression. A.** Mean squared error (MSE) are calculated between the predicted and observed 5hmC read counts in all gene bodies for 4 time points in "Forebrain Organoid". **B.** MSE are calculated between the predicted and observed gene expression for 4 time points in "Forebrain Organoid". **C.** MSE are calculated between the predicted and observed 5hmC read counts in all gene bodies for 17 human tissues in "Human Tissues". **D.** MSE are calculated between the predicted and observed gene expression for 17 human tissues in "Human Tissues".

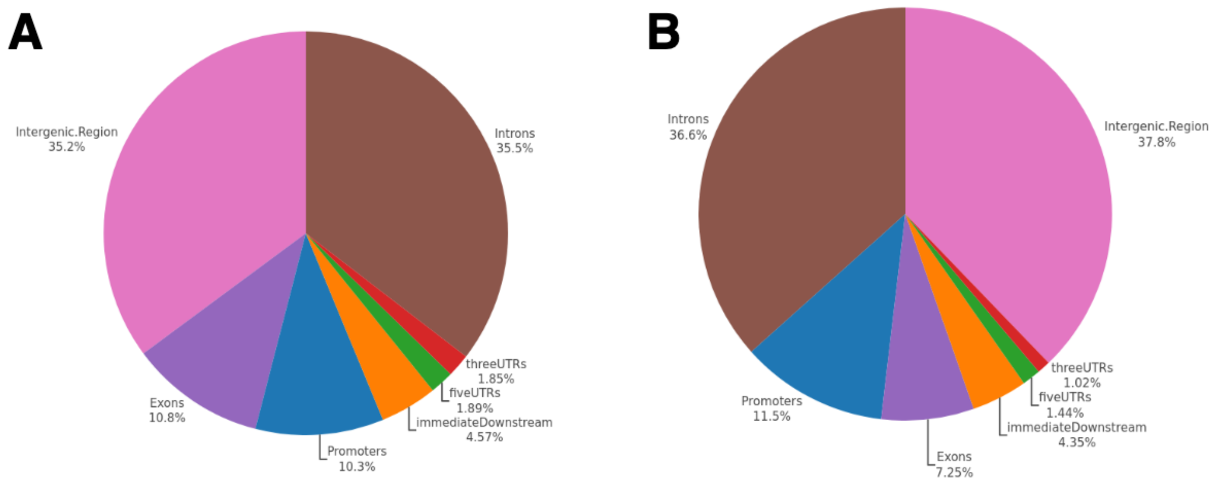


**Supplementary Figure S4. Comparing the distribution of DhMRs in the training set of "Kentucky AD" to genome-wide *de novo* DhMRs across different genomic features. A.** The distribution of DhMRs in the training set of "Kentucky AD" across different genomic features. **B.** The distribution of genome-wide *de novo* DhMRs across different genomic features.

### Supplementary Tables

| Histone Mark | ChIP-seq Data Source |
|---|---|
| H3K4me1 | BI.Brain_Angular_Gyrus.H3K4me1.112.filt.tagAlign.gz |
| H3K27ac | BI.Brain_Angular_Gyrus.H3K27ac.112.filt.tagAlign.gz |
| H3K9ac | BI.Brain_Angular_Gyrus.H3K9ac.112.filt.tagAlign.gz |
| H3K4me3 | BI.Brain_Angular_Gyrus.H3K4me3.112.filt.tagAlign.gz |
| H3K36me3 | BI.Brain_Angular_Gyrus.H3K36me3.112.filt.tagAlign.gz |
| H3K27me3 | BI.Brain_Angular_Gyrus.H3K27me3.149.filt.tagAlign.gz |
| H3K9me3 | BI.Brain_Angular_Gyrus.H3K9me3.112.filt.tagAlign.gz |

**Supplementary Table S1. Source of ChIP-seq data in "Brain Angular Gyrus" from Roadmap Epigenomics for exploring the distribution pattern of histone modification in the neighborhoods of EB 5hmC peaks from "Forebrain Organoid".**

| Histone Mark | ChIP-seq Data Source |
| --- | --- |
| H3K4me1 | BI.Brain_Angular_Gyrus.H3K4me1.112.filt.tagAlign.gz; BI.Brain_Angular_Gyrus.H3K4me1.149.filt.tagAlign.gz; BI.Brain_Anterior_Caudate.H3K4me1.112.filt.tagAlign.gz; BI.Brain_Anterior_Caudate.H3K4me1.149.filt.tagAlign.gz; BI.Brain_Cingulate_Gyrus.H3K4me1.112.filt.tagAlign.gz; BI.Brain_Cingulate_Gyrus.H3K4me1.149.filt.tagAlign.gz; BI.Brain_Hippocampus_Middle.H3K4me1.112.filt.tagAlign.gz; BI.Brain_Hippocampus_Middle.H3K4me1.149.filt.tagAlign.gz; BI.Brain_Hippocampus_Middle.H3K4me1.150.filt.tagAlign.gz; BI.Brain_Inferior_Temporal_Lobe.H3K4me1.112.filt.tagAlign.gz; BI.Brain_Inferior_Temporal_Lobe.H3K4me1.149.filt.tagAlign.gz; BI.Brain_Mid_Frontal_Lobe.H3K4me1.112.filt.tagAlign.gz; BI.Brain_Mid_Frontal_Lobe.H3K4me1.149.filt.tagAlign.gz; BI.Brain_Substantia_Nigra.H3K4me1.112.filt.tagAlign.gz; BI.Brain_Substantia_Nigra.H3K4me1.149.filt.tagAlign.gz; BI.Fetal_Brain.H3K4me1.UW_H22676.filt.tagAlign.gz; UCSF-UBC.Brain_Germinal_Matrix.H3K4me1.HuFGM01.filt.tagAlign.gz; UCSF-UBC.Brain_Germinal_Matrix.H3K4me1.HuFGM02.filt.tagAlign.gz; UCSF-UBC.Fetal_Brain.H3K4me1.HuFNSC01.filt.tagAlign.gz; UCSF-UBC.Fetal_Brain.H3K4me1.HuFNSC02.filt.tagAlign.gz |
| H3K27ac | BI.Brain_Angular_Gyrus.H3K27ac.112.filt.tagAlign.gz; BI.Brain_Angular_Gyrus.H3K27ac.149.filt.tagAlign.gz; BI.Brain_Anterior_Caudate.H3K27ac.112.filt.tagAlign.gz; BI.Brain_Anterior_Caudate.H3K27ac.149.filt.tagAlign.gz; BI.Brain_Cingulate_Gyrus.H3K27ac.112.filt.tagAlign.gz; BI.Brain_Cingulate_Gyrus.H3K27ac.149.filt.tagAlign.gz; BI.Brain_Hippocampus_Middle.H3K27ac.112.filt.tagAlign.gz; BI.Brain_Hippocampus_Middle.H3K27ac.149.filt.tagAlign.gz; BI.Brain_Hippocampus_Middle.H3K27ac.150.filt.tagAlign.gz; BI.Brain_Inferior_Temporal_Lobe.H3K27ac.112.filt.tagAlign.gz; BI.Brain_Inferior_Temporal_Lobe.H3K27ac.149.filt.tagAlign.gz; BI.Brain_Mid_Frontal_Lobe.H3K27ac.112.filt.tagAlign.gz; BI.Brain_Mid_Frontal_Lobe.H3K27ac.149.filt.tagAlign.gz; BI.Brain_Substantia_Nigra.H3K27ac.149.DNA_Lib_1847.filt.tagAlign.gz; BI.Brain_Substantia_Nigra.H3K27ac.149.filt.tagAlign.gz |
| H3K9ac | BI.Brain_Angular_Gyrus.H3K9ac.112.filt.tagAlign.gz; BI.Brain_Anterior_Caudate.H3K9ac.112.filt.tagAlign.gz; BI.Brain_Cingulate_Gyrus.H3K9ac.112.filt.tagAlign.gz; BI.Brain_Hippocampus_Middle.H3K9ac.112.filt.tagAlign.gz; BI.Brain_Inferior_Temporal_Lobe.H3K9ac.112.filt.tagAlign.gz; BI.Brain_Mid_Frontal_Lobe.H3K9ac.112.filt.tagAlign.gz; BI.Brain_Substantia_Nigra.H3K9ac.112.filt.tagAlign.gz; UCSF-UBC.Fetal_Brain.H3K9ac.HuFNSC-T.filt.tagAlign.gz |
| H3K4me3 | BI.Brain_Angular_Gyrus.H3K4me3.112.filt.tagAlign.gz; BI.Brain_Angular_Gyrus.H3K4me3.149.filt.tagAlign.gz; BI.Brain_Anterior_Caudate.H3K4me3.112.filt.tagAlign.gz; BI.Brain_Anterior_Caudate.H3K4me3.149.filt.tagAlign.gz; |

| | |
|---|---|
| | BI.Brain_Cingulate_Gyrus.H3K4me3.112.filt.tagAlign.gz;<br>BI.Brain_Cingulate_Gyrus.H3K4me3.149.filt.tagAlign.gz;<br>BI.Brain_Hippocampus_Middle.H3K4me3.112.filt.tagAlign.gz;<br>BI.Brain_Hippocampus_Middle.H3K4me3.149.filt.tagAlign.gz;<br>BI.Brain_Hippocampus_Middle.H3K4me3.150.filt.tagAlign.gz;<br>BI.Brain_Inferior_Temporal_Lobe.H3K4me3.112.filt.tagAlign.gz;<br>BI.Brain_Inferior_Temporal_Lobe.H3K4me3.149.filt.tagAlign.gz;<br>BI.Brain_Mid_Frontal_Lobe.H3K4me3.112.filt.tagAlign.gz;<br>BI.Brain_Mid_Frontal_Lobe.H3K4me3.149.filt.tagAlign.gz;<br>BI.Brain_Substantia_Nigra.H3K4me3.112.filt.tagAlign.gz;<br>BI.Brain_Substantia_Nigra.H3K4me3.149.filt.tagAlign.gz;<br>BI.Fetal_Brain.H3K4me3.UW_H-22510.filt.tagAlign.gz;<br>UCSF-UBC.Brain_Germinal_Matrix.H3K4me3.HuFGM01.filt.tagAlign.gz;<br>UCSF-UBC.Brain_Germinal_Matrix.H3K4me3.HuFGM02.filt.tagAlign.gz;<br>UCSF-UBC.Fetal_Brain.H3K4me3.HuFNSC-T.filt.tagAlign.gz;<br>UCSF-UBC.Fetal_Brain.H3K4me3.HuFNSC01.filt.tagAlign.gz;<br>UCSF-UBC.Fetal_Brain.H3K4me3.HuFNSC02.filt.tagAlign.gz |
| H3K36me3 | BI.Brain_Angular_Gyrus.H3K36me3.112.filt.tagAlign.gz;<br>BI.Brain_Angular_Gyrus.H3K36me3.149.filt.tagAlign.gz;<br>BI.Brain_Anterior_Caudate.H3K36me3.112.filt.tagAlign.gz;<br>BI.Brain_Anterior_Caudate.H3K36me3.149.filt.tagAlign.gz;<br>BI.Brain_Cingulate_Gyrus.H3K36me3.112.filt.tagAlign.gz;<br>BI.Brain_Cingulate_Gyrus.H3K36me3.149.filt.tagAlign.gz;<br>BI.Brain_Hippocampus_Middle.H3K36me3.112.filt.tagAlign.gz;<br>BI.Brain_Hippocampus_Middle.H3K36me3.149.filt.tagAlign.gz;<br>BI.Brain_Hippocampus_Middle.H3K36me3.150.filt.tagAlign.gz;<br>BI.Brain_Inferior_Temporal_Lobe.H3K36me3.112.filt.tagAlign.gz;<br>BI.Brain_Inferior_Temporal_Lobe.H3K36me3.149.filt.tagAlign.gz;<br>BI.Brain_Mid_Frontal_Lobe.H3K36me3.112.filt.tagAlign.gz;<br>BI.Brain_Mid_Frontal_Lobe.H3K36me3.149.filt.tagAlign.gz;<br>BI.Brain_Substantia_Nigra.H3K36me3.112.filt.tagAlign.gz;<br>BI.Brain_Substantia_Nigra.H3K36me3.149.filt.tagAlign.gz;<br>BI.Fetal_Brain.H3K36me3.UW_H-22510.filt.tagAlign.gz;<br>UCSF-UBC.Brain_Germinal_Matrix.H3K36me3.HuFGM01.filt.tagAlign.gz;<br>UCSF-UBC.Brain_Germinal_Matrix.H3K36me3.HuFGM02.filt.tagAlign.gz;<br>UCSF-UBC.Fetal_Brain.H3K36me3.HuFNSC01.filt.tagAlign.gz;<br>UCSF-UBC.Fetal_Brain.H3K36me3.HuFNSC02.filt.tagAlign.gz |
| H3K27me3 | BI.Brain_Angular_Gyrus.H3K27me3.149.filt.tagAlign.gz;<br>BI.Brain_Anterior_Caudate.H3K27me3.112.filt.tagAlign.gz;<br>BI.Brain_Anterior_Caudate.H3K27me3.149.filt.tagAlign.gz;<br>BI.Brain_Cingulate_Gyrus.H3K27me3.149.filt.tagAlign.gz;<br>BI.Brain_Hippocampus_Middle.H3K27me3.112.filt.tagAlign.gz;<br>BI.Brain_Hippocampus_Middle.H3K27me3.149.filt.tagAlign.gz;<br>BI.Brain_Hippocampus_Middle.H3K27me3.150.filt.tagAlign.gz;<br>BI.Brain_Inferior_Temporal_Lobe.H3K27me3.112.filt.tagAlign.gz;<br>BI.Brain_Inferior_Temporal_Lobe.H3K27me3.149.filt.tagAlign.gz;<br>BI.Brain_Mid_Frontal_Lobe.H3K27me3.149.filt.tagAlign.gz; |

| | |
|---|---|
| | BI.Brain_Substantia_Nigra.H3K27me3.112.filt.tagAlign.gz;<br>BI.Brain_Substantia_Nigra.H3K27me3.149.filt.tagAlign.gz;<br>BI.Fetal_Brain.H3K27me3.UW_H-22510.filt.tagAlign.gz;<br>BI.Fetal_Brain.H3K27me3.UW_H22676.filt.tagAlign.gz;<br>UCSF-UBC.Brain_Germinal_Matrix.H3K27me3.HuFGM01.filt.tagAlign.gz;<br>UCSF-UBC.Brain_Germinal_Matrix.H3K27me3.HuFGM02.filt.tagAlign.gz;<br>UCSF-UBC.Fetal_Brain.H3K27me3.HuFNSC-T.filt.tagAlign.gz;<br>UCSF-UBC.Fetal_Brain.H3K27me3.HuFNSC01.filt.tagAlign.gz;<br>UCSF-UBC.Fetal_Brain.H3K27me3.HuFNSC02.filt.tagAlign.gz |
| H3K9me3 | BI.Brain_Angular_Gyrus.H3K9me3.112.filt.tagAlign.gz;<br>BI.Brain_Angular_Gyrus.H3K9me3.149.filt.tagAlign.gz;<br>BI.Brain_Anterior_Caudate.H3K9me3.112.filt.tagAlign.gz;<br>BI.Brain_Anterior_Caudate.H3K9me3.149.filt.tagAlign.gz;<br>BI.Brain_Cingulate_Gyrus.H3K9me3.112.filt.tagAlign.gz;<br>BI.Brain_Cingulate_Gyrus.H3K9me3.149.filt.tagAlign.gz;<br>BI.Brain_Hippocampus_Middle.H3K9me3.112.filt.tagAlign.gz;<br>BI.Brain_Hippocampus_Middle.H3K9me3.149.filt.tagAlign.gz;<br>BI.Brain_Hippocampus_Middle.H3K9me3.150.filt.tagAlign.gz;<br>BI.Brain_Inferior_Temporal_Lobe.H3K9me3.112.filt.tagAlign.gz;<br>BI.Brain_Inferior_Temporal_Lobe.H3K9me3.149.filt.tagAlign.gz;<br>BI.Brain_Mid_Frontal_Lobe.H3K9me3.112.filt.tagAlign.gz;<br>BI.Brain_Mid_Frontal_Lobe.H3K9me3.149.filt.tagAlign.gz;<br>BI.Brain_Substantia_Nigra.H3K9me3.112.filt.tagAlign.gz;<br>BI.Brain_Substantia_Nigra.H3K9me3.149.filt.tagAlign.gz;<br>BI.Fetal_Brain.H3K9me3.UW_H-22510.filt.tagAlign.gz;<br>BI.Fetal_Brain.H3K9me3.UW_H22676.filt.tagAlign.gz;<br>UCSF-UBC.Brain_Germinal_Matrix.H3K9me3.HuFGM01.filt.tagAlign.gz;<br>UCSF-UBC.Brain_Germinal_Matrix.H3K9me3.HuFGM02.filt.tagAlign.gz;<br>UCSF-UBC.Fetal_Brain.H3K9me3.HuFNSC01.filt.tagAlign.gz;<br>UCSF-UBC.Fetal_Brain.H3K9me3.HuFNSC02.filt.tagAlign.gz |

**Supplementary Table S2. Source of ChIP-seq data in all brain regions from Roadmap Epigenomics for evaluating the predictive power of seven histone marks and being used by Deep5hmC as histone features (H3K4me1 and H3K4me3) for evaluating 4 developmental stages in "Forebrain Organoid".**

| Tissue | Histone mark | ChIP-seq Data Source |
|---|---|---|
| Bladder | H3K4me1 | UCSD.Bladder.H3K4me1.STL003.filt.tagAlign.gz |
| | H3K4me3 | ENCODE_ENCFF009JNQ_H3K4me3_Bladder.bed |
| Brain | H3K4me1 | BI.Brain_Angular_Gyrus.H3K4me1.112.filt.tagAlign.gz;<br>BI.Brain_Angular_Gyrus.H3K4me1.149.filt.tagAlign.gz;<br>BI.Brain_Anterior_Caudate.H3K4me1.112.filt.tagAlign.gz;<br>BI.Brain_Anterior_Caudate.H3K4me1.149.filt.tagAlign.gz;<br>BI.Brain_Cingulate_Gyrus.H3K4me1.112.filt.tagAlign.gz;<br>BI.Brain_Cingulate_Gyrus.H3K4me1.149.filt.tagAlign.gz;<br>BI.Brain_Hippocampus_Middle.H3K4me1.112.filt.tagAlign.gz;<br>BI.Brain_Hippocampus_Middle.H3K4me1.149.filt.tagAlign.gz;<br>BI.Brain_Hippocampus_Middle.H3K4me1.150.filt.tagAlign.gz;<br>BI.Brain_Inferior_Temporal_Lobe.H3K4me1.112.filt.tagAlign.gz;<br>BI.Brain_Inferior_Temporal_Lobe.H3K4me1.149.filt.tagAlign.gz;<br>BI.Brain_Mid_Frontal_Lobe.H3K4me1.112.filt.tagAlign.gz;<br>BI.Brain_Mid_Frontal_Lobe.H3K4me1.149.filt.tagAlign.gz;<br>BI.Brain_Substantia_Nigra.H3K4me1.112.filt.tagAlign.gz;<br>BI.Brain_Substantia_Nigra.H3K4me1.149.filt.tagAlign.gz;<br>BI.Fetal_Brain.H3K4me1.UW_H22676.filt.tagAlign.gz;<br>UCSF-UBC.Brain_Germinal_Matrix.H3K4me1.HuFGM01.filt.tagAlign.gz;<br>UCSF-UBC.Brain_Germinal_Matrix.H3K4me1.HuFGM02.filt.tagAlign.gz;<br>UCSF-UBC.Fetal_Brain.H3K4me1.HuFNSC01.filt.tagAlign.gz;<br>UCSF-UBC.Fetal_Brain.H3K4me1.HuFNSC02.filt.tagAlign.gz |
| | H3K4me3 | BI.Brain_Angular_Gyrus.H3K4me3.112.filt.tagAlign.gz;<br>BI.Brain_Angular_Gyrus.H3K4me3.149.filt.tagAlign.gz;<br>BI.Brain_Anterior_Caudate.H3K4me3.112.filt.tagAlign.gz;<br>BI.Brain_Anterior_Caudate.H3K4me3.149.filt.tagAlign.gz;<br>BI.Brain_Cingulate_Gyrus.H3K4me3.112.filt.tagAlign.gz;<br>BI.Brain_Cingulate_Gyrus.H3K4me3.149.filt.tagAlign.gz;<br>BI.Brain_Hippocampus_Middle.H3K4me3.112.filt.tagAlign.gz;<br>BI.Brain_Hippocampus_Middle.H3K4me3.149.filt.tagAlign.gz;<br>BI.Brain_Hippocampus_Middle.H3K4me3.150.filt.tagAlign.gz;<br>BI.Brain_Inferior_Temporal_Lobe.H3K4me3.112.filt.tagAlign.gz;<br>BI.Brain_Inferior_Temporal_Lobe.H3K4me3.149.filt.tagAlign.gz;<br>BI.Brain_Mid_Frontal_Lobe.H3K4me3.112.filt.tagAlign.gz;<br>BI.Brain_Mid_Frontal_Lobe.H3K4me3.149.filt.tagAlign.gz;<br>BI.Brain_Substantia_Nigra.H3K4me3.112.filt.tagAlign.gz;<br>BI.Brain_Substantia_Nigra.H3K4me3.149.filt.tagAlign.gz;<br>BI.Fetal_Brain.H3K4me3.UW_H-22510.filt.tagAlign.gz;<br>UCSF-UBC.Brain_Germinal_Matrix.H3K4me3.HuFGM01.filt.tagAlign.gz;<br>UCSF-UBC.Brain_Germinal_Matrix.H3K4me3.HuFGM02.filt.tagAlign.gz;<br>UCSF-UBC.Fetal_Brain.H3K4me3.HuFNSC-T.filt.tagAlign.gz;<br>UCSF-UBC.Fetal_Brain.H3K4me3.HuFNSC01.filt.tagAlign.gz;<br>UCSF-UBC.Fetal_Brain.H3K4me3.HuFNSC02.filt.tagAlign.gz |
| Breast | H3K4me1 | UCSF-<br>UBC.Breast_Fibroblast_Primary_Cells.H3K4me1.RM071.filt.tagAlign.gz;<br>UCSF-<br>UBC.Breast_Luminal_Epithelial_Cells.H3K4me1.RM080.filt.tagAlign.gz;<br>UCSF-UBC.Breast_Myoepithelial_Cells.H3K4me1.RM066.filt.tagAlign.gz;<br>UCSF-UBC.Breast_Myoepithelial_Cells.H3K4me1.RM080.filt.tagAlign.gz; |

| | | |
|---|---|---|
| | | UCSF-UBC.Breast_vHMEC.H3K4me1.RM035.HS1994.filt.tagAlign.gz; UCSF-UBC.Breast_vHMEC.H3K4me1.RM035.HS2618.filt.tagAlign.gz |
| | H3K4me3 | UCSF-UBC.Breast_Fibroblast_Primary_Cells.H3K4me3.RM071.filt.tagAlign.gz; UCSF-UBC.Breast_Myoepithelial_Cells.H3K4me3.RM066.filt.tagAlign.gz; UCSF-UBC.Breast_Myoepithelial_Cells.H3K4me3.RM080.filt.tagAlign.gz; UCSF-UBC.Breast_vHMEC.H3K4me3.RM035.HS2615.filt.tagAlign.gz |
| Heart | H3K4me1 | BI.Fetal_Heart.H3K4me1.H-23524.filt.tagAlign.gz; BI.Fetal_Heart.H3K4me1.UW_H23914.filt.tagAlign.gz |
| | H3K4me3 | BI.Fetal_Heart.H3K4me3.UW_H23914.filt.tagAlign.gz |
| Kidney | H3K4me1 | BI.Adult_Kidney.H3K4me1.153.filt.tagAlign.gz; BI.Adult_Kidney.H3K4me1.27.filt.tagAlign.gz; BI.Fetal_Kidney.H3K4me1.UW_H-22676.filt.tagAlign.gz |
| | H3K4me3 | BI.Adult_Kidney.H3K4me3.153.filt.tagAlign.gz; BI.Adult_Kidney.H3K4me3.27.filt.tagAlign.gz; BI.Fetal_Kidney.H3K4me3.UW_H-22676.filt.tagAlign.gz |
| Liver | H3K4me1 | BI.Adult_Liver.H3K4me1.3.filt.tagAlign.gz; BI.Adult_Liver.H3K4me1.4.filt.tagAlign.gz; BI.Adult_Liver.H3K4me1.5.filt.tagAlign.gz; UCSD.Adult_Liver.H3K4me1.STL011.filt.tagAlign.gz |
| | H3K4me3 | BI.Adult_Liver.H3K4me3.3.filt.tagAlign.gz; BI.Adult_Liver.H3K4me3.4.filt.tagAlign.gz; BI.Adult_Liver.H3K4me3.5.filt.tagAlign.gz; UCSD.Adult_Liver.H3K4me3.STL011.filt.tagAlign.gz |
| Lung | H3K4me1 | BI.Fetal_Lung.H3K4me1.UW_H-22727.filt.tagAlign.gz; BI.Fetal_Lung.H3K4me1.UW_H22772.filt.tagAlign.gz; BI.Fetal_Lung.H3K4me1.UW_H23266.filt.tagAlign.gz; UCSD.Lung.H3K4me1.STL001.filt.tagAlign.gz; UCSD.Lung.H3K4me1.STL002.filt.tagAlign.gz |
| | H3K4me3 | BI.Fetal_Lung.H3K4me3.UW_H-22676.filt.tagAlign.gz; BI.Fetal_Lung.H3K4me3.UW_H-22727.filt.tagAlign.gz; UCSD.Lung.H3K4me3.STL002.filt.tagAlign.gz |
| Marrow | H3K4me1 | BI.Bone_Marrow_Derived_Mesenchymal_Stem_Cell_Cultured_Cells.H3K4me1.57.filt.tagAlign.gz; BI.Bone_Marrow_Derived_Mesenchymal_Stem_Cell_Cultured_Cells.H3K4me1.58.filt.tagAlign.gz; BI.Bone_Marrow_Derived_Mesenchymal_Stem_Cell_Cultured_Cells.H3K4me1.59.filt.tagAlign.gz; BI.Bone_Marrow_Derived_Mesenchymal_Stem_Cell_Cultured_Cells.H3K4me1.60.filt.tagAlign.gz; BI.Chondrocytes_from_Bone_Marrow_Derived_Mesenchymal_Stem_Cell_Cultured_Cells.H3K4me1.57.filt.tagAlign.gz; BI.Chondrocytes_from_Bone_Marrow_Derived_Mesenchymal_Stem_Cell_Cultured_Cells.H3K4me1.58.filt.tagAlign.gz; BI.Chondrocytes_from_Bone_Marrow_Derived_Mesenchymal_Stem_Cell_Cultured_Cells.H3K4me1.59.filt.tagAlign.gz; BI.Chondrocytes_from_Bone_Marrow_Derived_Mesenchymal_Stem_Cell_Cultured_Cells.H3K4me1.60.filt.tagAlign.gz |
| | H3K4me3 | BI.Bone_Marrow_Derived_Mesenchymal_Stem_Cell_Cultured_Cells.H3K4me3.57.filt.tagAlign.gz; |

| | | |
|---|---|---|
| | | BI.Bone_Marrow_Derived_Mesenchymal_Stem_Cell_Cultured_Cells.H3K4me3.58.filt.tagAlign.gz; BI.Bone_Marrow_Derived_Mesenchymal_Stem_Cell_Cultured_Cells.H3K4me3.59.filt.tagAlign.gz; BI.Bone_Marrow_Derived_Mesenchymal_Stem_Cell_Cultured_Cells.H3K4me3.60.filt.tagAlign.gz; BI.Chondrocytes_from_Bone_Marrow_Derived_Mesenchymal_Stem_Cell_Cultured_Cells.H3K4me3.57.filt.tagAlign.gz; BI.Chondrocytes_from_Bone_Marrow_Derived_Mesenchymal_Stem_Cell_Cultured_Cells.H3K4me3.58.filt.tagAlign.gz; BI.Chondrocytes_from_Bone_Marrow_Derived_Mesenchymal_Stem_Cell_Cultured_Cells.H3K4me3.59.filt.tagAlign.gz; BI.Chondrocytes_from_Bone_Marrow_Derived_Mesenchymal_Stem_Cell_Cultured_Cells.H3K4me3.60.filt.tagAlign.gz |
| Ovary | H3K4me1 | UCSD.Ovary.H3K4me1.STL002.filt.tagAlign.gz |
| | H3K4me3 | UCSD.Ovary.H3K4me3.STL002.filt.tagAlign.gz |
| Pancreas | H3K4me1 | UCSD.Pancreas.H3K4me1.STL002.filt.tagAlign.gz; UCSD.Pancreas.H3K4me1.STL003.filt.tagAlign.gz |
| | H3K4me3 | UCSD.Pancreas.H3K4me3.STL003.filt.tagAlign.gz |
| Placenta | H3K4me1 | UCSF-UBC.Placenta_Amnion.H3K4me1.CTL02.filt.tagAlign.gz; UCSF-UBC.Placenta_Chorion_Smooth.H3K4me1.CTL02.filt.tagAlign.gz; UW.Fetal_Placenta.H3K4me1.H-24996.Histone.DS23027.filt.tagAlign.gz |
| | H3K4me3 | UCSF-UBC.Placenta_Amnion.H3K4me3.CTL02.filt.tagAlign.gz; UCSF-UBC.Placenta_Chorion_Smooth.H3K4me3.CTL02.filt.tagAlign.gz; UW.Fetal_Placenta.H3K4me3.H-24996.Histone.DS23300.filt.tagAlign.gz |
| Prostate | H3K4me1 | ENCODE_ENCFF061CPC_H3K4me1_Prostate.bed; ENCODE_ENCFF099KSH_H3K4me1_Prostate.bed; ENCODE_ENCFF162ACA_H3K4me1_Prostate.bed; ENCODE_ENCFF324PUS_H3K4me1_Prostate.bed |
| | H3K4me3 | ENCODE_ENCFF055CAO_H3K4me3_Prostate.bed; ENCODE_ENCFF369SLA_H3K4me3_Prostate.bed; ENCODE_ENCFF483OJJ_H3K4me3_Prostate.bed; ENCODE_ENCFF854ANC_H3K4me3_Prostate.bed |
| Colon (Sigmoid) | H3K4me1 | UCSD.Sigmoid_Colon.H3K4me1.STL001.filt.tagAlign.gz; UCSD.Sigmoid_Colon.H3K4me1.STL003.filt.tagAlign.gz |
| | H3K4me3 | UCSD.Sigmoid_Colon.H3K4me3.STL001.filt.tagAlign.gz; UCSD.Sigmoid_Colon.H3K4me3.STL003.filt.tagAlign.gz |
| Skin | H3K4me1 | ENCODE_ENCFF797BMX_H3K4me1_Skin.bed; UCSF-UBC.Penis_Foreskin_Fibroblast_Primary_Cells.H3K4me1.skin01.filt.tagAlign.gz; UCSF-UBC.Penis_Foreskin_Fibroblast_Primary_Cells.H3K4me1.skin02.filt.tagAlign.gz; UCSF-UBC.Penis_Foreskin_Fibroblast_Primary_Cells.H3K4me1.skin03.filt.tagAlign.gz; UCSF-UBC.Penis_Foreskin_Keratinocyte_Primary_Cells.H3K4me1.skin01.filt.tagAlign.gz; |

| | | |
|---|---|---|
| | | UCSF-UBC.Penis_Foreskin_Keratinocyte_Primary_Cells.H3K4me1.skin02.filt.tagAlign.gz;<br>UCSF-UBC.Penis_Foreskin_Keratinocyte_Primary_Cells.H3K4me1.skin03.filt.tagAlign.gz;<br>UCSF-UBC.Penis_Foreskin_Melanocyte_Primary_Cells.H3K4me1.skin01.filt.tagAlign.gz;<br>UCSF-UBC.Penis_Foreskin_Melanocyte_Primary_Cells.H3K4me1.skin02.filt.tagAlign.gz;<br>UCSF-UBC.Penis_Foreskin_Melanocyte_Primary_Cells.H3K4me1.skin03.filt.tagAlign.gz |
| | H3K4me3 | ENCODE_ENCFF258WJE_H3K4me3_Skin.bed;<br>UCSF-UBC.Penis_Foreskin_Fibroblast_Primary_Cells.H3K4me3.skin01.filt.tagAlign.gz;<br>UCSF-UBC.Penis_Foreskin_Fibroblast_Primary_Cells.H3K4me3.skin02.filt.tagAlign.gz;<br>UCSF-UBC.Penis_Foreskin_Fibroblast_Primary_Cells.H3K4me3.skin03.filt.tagAlign.gz;<br>UCSF-UBC.Penis_Foreskin_Keratinocyte_Primary_Cells.H3K4me3.skin01.filt.tagAlign.gz;<br>UCSF-UBC.Penis_Foreskin_Keratinocyte_Primary_Cells.H3K4me3.skin02.filt.tagAlign.gz;<br>UCSF-UBC.Penis_Foreskin_Keratinocyte_Primary_Cells.H3K4me3.skin03.filt.tagAlign.gz;<br>UCSF-UBC.Penis_Foreskin_Melanocyte_Primary_Cells.H3K4me3.skin01.filt.tagAlign.gz;<br>UCSF-UBC.Penis_Foreskin_Melanocyte_Primary_Cells.H3K4me3.skin02.filt.tagAlign.gz;<br>UCSF-UBC.Penis_Foreskin_Melanocyte_Primary_Cells.H3K4me3.skin03.filt.tagAlign.gz |
| Stomach | H3K4me1 | BI.Stomach_Mucosa.H3K4me1.157.filt.tagAlign.gz;<br>BI.Stomach_Smooth_Muscle.H3K4me1.28.filt.tagAlign.gz;<br>UW.Fetal_Stomach.H3K4me1.H-24776.Histone.DS22597.filt.tagAlign.gz |
| | H3K4me3 | BI.Stomach_Mucosa.H3K4me3.157.filt.tagAlign.gz;<br>BI.Stomach_Smooth_Muscle.H3K4me3.161.filt.tagAlign.gz;<br>BI.Stomach_Smooth_Muscle.H3K4me3.28.filt.tagAlign.gz;<br>UW.Fetal_Stomach.H3K4me3.H-24639.Histone.DS22598.filt.tagAlign.gz |

| | | |
|---|---|---|
| Testis | H3K4me1 | ENCODE_ENCFF020VQJ_H3K4me1_Testis.bed |
| | H3K4me3 | ENCODE_ENCFF007LNP_H3K4me3_Testis.bed; ENCODE_ENCFF796PVK_H3K4me3_Testis.bed |
| Colon (Transverse) | H3K4me1 | ENCODE_ENCFF002YUH_H3K4me1_Colon_Transverse.bed; ENCODE_ENCFF159ZCY_H3K4me1_Colon_Transverse.bed; ENCODE_ENCFF250ADB_H3K4me1_Colon_Transverse.bed; ENCODE_ENCFF530AJW_H3K4me1_Colon_Transverse.bed; ENCODE_ENCFF638YYL_H3K4me1_Colon_Transverse.bed; ENCODE_ENCFF804UEW_H3K4me1_Colon_Transverse.bed; ENCODE_ENCFF813QQE_H3K4me1_Colon_Transverse.bed; ENCODE_ENCFF996EQE_H3K4me1_Colon_Transverse.bed |
| | H3K4me3 | ENCODE_ENCFF128IEV_H3K4me3_Colon_Transverse.bed; ENCODE_ENCFF197YTB_H3K4me3_Colon_Transverse.bed; ENCODE_ENCFF276GOD_H3K4me3_Colon_Transverse.bed; ENCODE_ENCFF598QBZ_H3K4me3_Colon_Transverse.bed; ENCODE_ENCFF600EPC_H3K4me3_Colon_Transverse.bed; ENCODE_ENCFF614HSP_H3K4me3_Colon_Transverse.bed; ENCODE_ENCFF771GCJ_H3K4me3_Colon_Transverse.bed |

**Supplementary Table S3. Source of ChIP-seq data from ENCODE being used by Deep5hmC as histone features (H3K4me1 and H3K4me3) for evaluating 17 human tissues in "Human Tissues".**

| **Condition** | **Histone Mark** | **Gender** | **ChIP-seq Data Source** |
|---|---|---|---|
| Alzheimer's disease | H3K27ac | female | ENCFF167PHR |
| | H3K4me3 | female | ENCFF581GEK |
| Healthy control | H3K27ac | female | ENCFF372LKU |
| | H3K4me3 | female | ENCFF111DCY |

**Supplementary Table S4. Source of ChIP-seq data from ENCODE used by Deep5hmC as histone features (H3K27ac and H3K4me3) for predicting DhMRs in "Kentucky AD".**

| Data | Tissue | Number of Positive Peaks |
| --- | --- | --- |
| Brain Organoid | EB | 64458 |
| | D56 | 56036 |
| | D84 | 81032 |
| | D112 | 81050 |
| Human Tissues | Bladder | 98256 |
| | Brain | 29079 |
| | Breast | 46100 |
| | Heart | 75606 |
| | Kidney | 75959 |
| | Liver | 78819 |
| | Lung | 51466 |
| | Marrow | 12596 |
| | Ovary | 116948 |
| | Pancreas | 81555 |
| | Placenta | 137488 |
| | Prostate | 75271 |
| | Colon (Sigmoid) | 25428 |
| | Skin | 69322 |
| | Stomach | 28270 |
| | Testis | 25142 |
| | Colon (Transverse) | 32460 |
| Kentucky AD | - | 4330 |

**Supplementary Table S5. Summary of sample size for “Brain Organoid”, “Human Tissues” and “Kentucky AD” data.**
